# ApoE lipidation, not isoform, is the key modulator of Aβ interaction, uptake, and cytotoxicity

**DOI:** 10.1101/2025.11.19.685371

**Authors:** Kamila Nurmakova, Gregory-Neal W. Gomes, Zachary A. Levine

## Abstract

**Background:** Inherited variations in the Apolipoprotein E (*APOE*) gene are the largest genetic determinant for late-onset Alzheimer’s Disease, with the *APOEε4* allele conferring the highest risk. While *APOE* was shown to modulate amyloid beta (Aβ) pathology in a genotype-specific manner (*APOEε4*>*APOEε3*>*APOEε2*), it remains an important open question whether these differences are due directly to isoform-specific interactions between ApoE and Aβ or indirect effects on Aβ clearance. This is further complicated because ApoE exists in lipidated or unlipidated states, which influence its biochemical properties. To disentangle how single ApoE mutations confer vastly different effects on Aβ pathology, we investigated how both the ApoE isoform and lipidation modulate its interaction with inert and cytotoxic Aβ species, and its effect on Aβ uptake and cytotoxicity in human astrocytes.

**Methods:** We prepared Aβ in distinct aggregation states (monomers, oligomers, and fibrils) and ApoE in different lipidation states to characterize their size and affinity for one another using fluorescence correlation spectroscopy and fluorescence polarization, respectively. We then utilized flow cytometry and live cell imaging to quantify the uptake of Aβ in different aggregation states by primary and immortalized human astrocytes in the presence of different isoforms of lipidated or unlipidated ApoE.

**Results:** This study revealed that ApoE lipidation, not isoform, has the biggest impact on its interaction with Aβ, on the uptake of Aβ by astrocytes, and on Aβ-induced cytotoxicity. Specifically, unlipidated ApoE preferentially interacts with Aβ oligomers and fibrils, which substantially inhibits their uptake by astrocytes. Conversely, lipidated ApoE showed no interaction with Aβ oligomers and had a reduced ability to inhibit Aβ uptake.

**Conclusions:** Our observations provide important molecular details that suggest previously observed ApoE isoform-specific differences in AD risk are likely driven primarily by *in vivo* differences in ApoE lipidation, rather than biophysical differences between ApoE isoforms in their interaction with Aβ. We propose that the impaired lipidation of ApoE4 in the brain increases levels of unlipidated ApoE, which then bind toxic Aβ oligomers and reduce their clearance by astrocytes. These findings underscore the therapeutic potential of interventions aimed at increasing ApoE lipidation and decreasing interactions between toxic Aβ oligomers and ApoE.

## BACKGROUND

Alzheimer’s disease (AD) is the leading cause of dementia and is pathologically defined by the progressive accumulation of extracellular amyloid beta (Aβ) plaques and intracellular neurofibrillary tangles of hyperphosphorylated tau^1^. While previous research efforts heavily focused on the deposition of Aβ fibrillar plaques^2^, Aβ oligomers were recently recognized as central in AD pathology^3,4,5,6^. Unlike plaque burden, Aβ oligomer levels more accurately predict AD pathology and disease progression^7^. They initiate pathology through a variety of mechanisms, including the disruption of membrane integrity and the initiation of inflammation and cellular senescence^4^. In Familial Alzheimer’s Disease (FAD), genetic evidence strongly supports a direct link between Aβ and pathology through mutations to the amyloid precursor protein (*APP*), presenilin 1 (*PSEN1*), and presenilin 2 (*PSEN2*) that result in increased Aβ production and aggregation^8^. Despite strong Aβ-dependent genetic determinants of FAD, these cases account for only a minor fraction of all AD cases^9^. The vast majority involve sporadic late-onset AD (LOAD), where pathology is thought to progress through the defective clearance of Aβ rather than its overproduction^10^.

Among the many genes that impact Aβ clearance in the brain, Apolipoprotein E (*APOE*) has been identified as the strongest genetic risk factor for LOAD. ApoE is the major lipid carrier in the central nervous system and is predominantly expressed by astrocytes^11^. Humans possess three major alleles of *APOE* (*APOEε2*, *APOEε3*, and *APOEε4*), which differ in their risks of Alzheimer’s Disease^12^. Population genetics studies identify *APOEε4* as the strongest genetic risk factor for developing sporadic LOAD compared to the WT *APOEε3* allele^13,14,15^, whereas the *APOEε2* allele is protective against AD. The genotype-specific disease risks also correlate with changes to Aβ pathology. For example, levels of Aβ oligomers are highest in the brains of *APOEε4* carriers and lowest in *APOEε2* carriers^16^. Studies in transgenic mice also support these observations, showing that *APOEε4* mice have increased levels of soluble (monomers and oligomers) and insoluble Aβ along with impaired clearance, whereas *APOEε2* mice demonstrate lower Aβ levels and enhanced clearance^17,18,19^. Despite the established detrimental effects of *APOEε4* and protective effects of *APOEε2* alleles, there is no consensus regarding the mechanistic explanation for these observed differences in Aβ levels and clearance.

Previous studies have attempted to characterize the biophysical interactions between Aβ and ApoE^18,20,21^, and in particular have probed how ApoE modulates Aβ uptake in glial cells^20,22,23^. However, interpreting these studies can be particularly challenging or even disparate due to the many states that ApoE can adopt (isoform differences, lipidation, self-association, abundance, and stability). ApoE lipidation, for example, is a critical factor influencing its biological properties that directly modulates its stability and activity^24,25^. The degree of lipidation differs among ApoE isoforms, with ApoE4 forming more lipid-free and poorly-lipidated particles^26,27^. Given that lipids and Aβ share binding sites on ApoE^28^, lipidation of ApoE likely has a significant effect on its interaction with Aβ and its subsequent uptake by glial cells. Additionally, Aβ exists in different aggregation states (monomers, oligomers, and fibrils) that coexist along the aggregation pathway and differ in their structures and biological activity^29^.

Conflicting data regarding the role of ApoE in modulating Aβ clearance significantly limit the understanding of AD pathology and represent a hindrance to the development of targeted therapeutic strategies. Considering the individual and combined molecular complexity of Aβ and ApoE, there is a substantial need for systematic studies to decouple how ApoE isoform (ApoE2 vs. ApoE3 vs. ApoE4), lipidation (lipidated vs. unlipidated), and Aβ aggregation state (monomers vs. oligomers vs. fibrils) jointly influence their interaction and cellular uptake.

To address these questions, we selectively prepared Aβ in different aggregation states and investigated their interaction with both lipidated and unlipidated ApoE. Doing so enabled highly reproducible observations that provide a robust foundation for future comparative and therapeutic studies. We used fluorescence correlation spectroscopy (FCS) to characterize the sizes of ApoE and Aβ followed by fluorescence polarization (FP) to characterize the affinity of ApoE:Aβ complexes. Next, we looked at the uptake of Aβ monomers, oligomers, and fibrils by human cortical astrocytes, which are critical regulators of Aβ metabolism^30,31^. Preparing stabilized Aβ in distinct aggregation states allowed us to independently measure the clearance of individual Aβ species such as toxic Aβ oligomers, which are often difficult to distinguish from non-toxic Aβ species.

Surprisingly, we find that when ApoE isoforms were equally lipidated, their interaction with Aβ or their ability to modulate Aβ uptake and cytotoxicity did not extensively differ. Conversely, the lipidation of ApoE, regardless of isoform, was the strongest determinant of its affinity with Aβ oligomers, Aβ oligomer uptake, and Aβ oligomer cytotoxicity. Given that ApoE isoforms are differentially lipidated *in vivo*^26,27^, our study suggests that ApoE lipidation is the primary driver of isoform-dependent differences in AD pathology. As a result, we propose that the increased *in vivo* lipidation of ApoE2 relative to ApoE3 would provide protection from toxic Aβ oligomers. Alternatively, decreased lipidation of ApoE4 likely increases the interaction with toxic Aβ oligomers and inhibits astrocytic uptake, resulting in increased extracellular concentrations of toxic Aβ oligomers. This scenario also suggests that the protective effects of ApoE2 and pathogenic effects of ApoE4 are not primarily determined by isoform-specific interactions with Aβ, but rather by isoform-specific differences in ApoE lipidation that in turn affect the extracellular concentration of toxic Aβ oligomers. These observations also clarify that direct interactions between Aβ and ApoE take place in distinct states that must be deconvoluted in systematic studies. Doing so not only improves the reliability and reproducibility of cellular and biochemical experiments, but also informs the mechanistic crosstalk between Aβ and ApoE in AD pathology in order to vastly simplify therapeutic strategies to combat the root causes of neurodegeneration.

## RESULTS

### FCS confirms distinct and reproducible Aβ monomers, oligomers, and fibrils

Distinguishing between the aggregation states of Aβ is crucial when examining their interaction with other proteins and their effects on cells due to the differences in their biochemical and biological properties, such as size, solubility, and cytotoxicity. Therefore, we separately prepared Aβ monomers, oligomers, and fibrils and characterized their biophysical properties to normalize for their aggregation state and ensure reproducibility for the following experiments. We recombinantly expressed Aβ42 in *E. coli* using the protocol outlined by the Linse lab^32^ and assessed peptide quality via a thioflavin T (ThT) aggregation assay, which reports on fibrillar content in solution^33^. A peptide solution of Aβ at 2.5 μM, a concentration commonly used in the ThT assays, showed a low initial fluorescence in PBS, an extended lag time of 59 ± 5 minutes, and a reproducible t_1/2_ of 79 ± 8 minutes, which is indicative of an aggregate-free monomeric starting material (Fig. S1C). Next, we synthesized stable Aβ oligomers following the protocol by Barghorn *et al.*^34^. Pre-formed oligomers enabled us to circumvent the difficulties associated with sample heterogeneity or the transient nature of the oligomers. We also prepared Aβ fibrils by aggregating Aβ at 100 μM concentration in 10 mM HCl as previously published^35^.

We then used FCS to characterize the sizes of Aβ in different aggregation states. FCS is a statistical technique that measures the diffusion characteristics of fluorescently labeled molecules by observing the correlated fluorescence intensity fluctuations as molecules pass through a small confocal volume^36^, which allows us to determine the hydrodynamic radii (R_H_) of Aβ monomers, oligomers, and fibrils. First, we looked at Alexa Fluor 594-labeled monomeric Aβ. The autocorrelation curve of Aβ monomers corresponded to a significantly smaller diffusion time when compared to Aβ oligomers and fibrils (Fig. 1A). We fit the FCS autocorrelation curve with a one-component (single-species) diffusion model (as described in “Materials and Methods”), which resulted in a diffusion time corresponding to a hydrodynamic radius (R_H_) of 1.76 ± 0.05 nm, consistent with previous reports for monomeric Aβ42^37^ (Fig. 1B). Importantly, we also confirmed that even at 1 μM, which is the concentration used in the uptake studies, Aβ remained monomeric in cell media at 37 °C over the 24 hour period (Fig. S1D, E).

**Figure 1.**
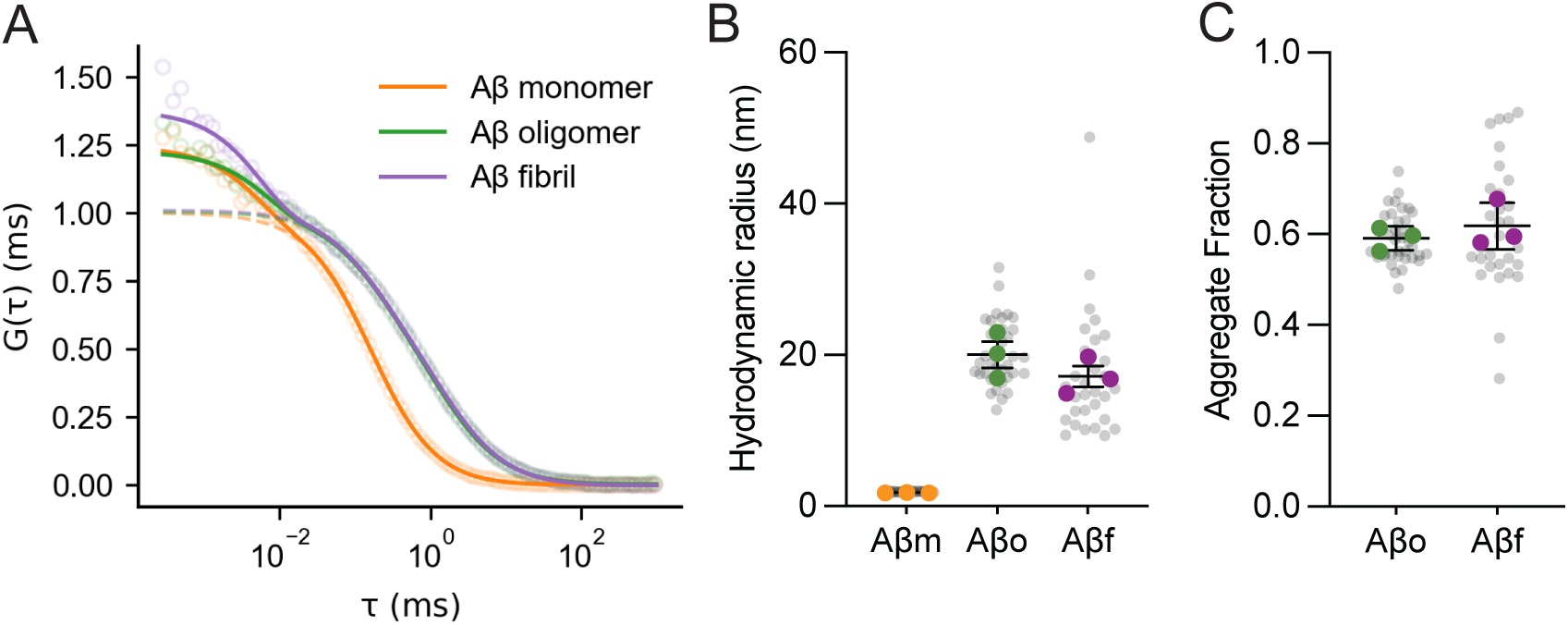
Characterization of Aβ monomers, oligomers, and fibrils via FCS. **(A)** Representative experimental (markers) and fitted (lines) autocorrelation curves for Aβ monomers (1 nM), Aβ oligomers (3 µM total Aβ, with 15 nM labeled Aβ incorporated), and sonicated Aβ fibrils (3 µM total Aβ, 1.5 nM labeled Aβ incorporated). The full fits, including a triplet component, are shown with solid lines. All autocorrelation curves and fits are normalized to the particle number N. **(B)** Hydrodynamic radii and **(C)** fractions of aggregates, as derived from autocorrelation curves. Independent Aβ monomer, oligomer, and fibril preparations are shown in orange, green, and purple markers, along with mean and SEM (n=3). Gray markers indicate individual 30-second measurements within the same preparation (10 per sample).

To prepare fluorescently labeled Aβ oligomers and fibrils, we incorporated a small amount of Alexa Fluor 594-labeled Aβ monomer into the high excess of unlabeled Aβ monomer during the Aβ oligomer and fibril preparation steps (outlined in “Materials and Methods”). We optimized the labeling ratio to minimize the amount of label per aggregate, while also maintaining a high enough ratio for a detectable fluorescence intensity signal at each concentration. In contrast to Aβ monomers, Alexa Fluor 594-labeled Aβ oligomers required a two-component fit, which indicated the presence of multiple diffusing species. Since Aβ oligomers exist in equilibrium with Aβ monomers in solution^38^, we fixed one of the diffusion components to the monomeric Aβ diffusion time. The second, slower-diffusing component corresponded to a large Aβ oligomer population with an R_H_ of 20 ± 3 nm (Fig. 1B). These oligomers constituted 59 ± 2% of total fluorescently labeled species in solution (Fig. 1C).

Due to their larger size, Aβ fibrils were unable to diffuse through the confocal volume. To produce smaller diffusing fragments, we sonicated Aβ fibrils for 10 minutes. Sonicated Aβ fibrils required a two-component diffusion fit with one component fixed to the monomer diffusion time, similar to Aβ oligomers. Sonicated Aβ fibrils had a mean R_H_ of 17 ± 2 nm (Fig. 1B), which was comparable to Aβ oligomers. Thus, our sonicated Aβ fibrils can be thought of as fibrillar oligomers that are distinct from non-fibrillar oligomers. The fraction of these slow-diffusing sonicated Aβ fibrils was comparable to Aβ oligomers (62 ± 5%) (Fig. 1C). We also assessed the reproducibility of individual preparations (Fig 1B, C, green and purple markers), and the heterogeneity within Aβ aggregate preparations using a previously described approach of sampling multiple short (30 s) correlation times and fitting these individual autocorrelation curves^39^ (Fig 1B, C, grey markers). Three independent Aβ oligomer and fibril preparations resulted in both highly reproducible R_H_ and slow component fractions. However, we found that Aβ fibrils exhibited a larger distribution of sizes within the same preparation (CV = 43 ± 25%) compared to Aβ oligomers (CV = 17 ± 2%), indicating a higher heterogeneity of fibrils.

### Aβ uptake by human astrocytes is highly dependent on its aggregation state

Uptake of Aβ by astrocytes is a critical process in Aβ metabolism^40^, however, it is unclear if the rate of uptake differs between the distinct aggregation states of Aβ. To answer this question we employed live-cell imaging to characterize the uptake of Aβ by primary and immortalized human astrocytes, exposing them to equal concentrations of Aβ monomers, oligomers, and fibrils (containing the same amounts of fluorescent label) for 24 hours. We then combined fluorescence microscopy and flow cytometry to quantitatively assess the cellular association and uptake of each Aβ aggregation state.

Live cell imaging revealed significant cellular surface binding and uptake of Aβ oligomers and fibrils into endosomes, characterized by intracellular puncta (Fig. 2A). In contrast, Aβ monomers did not demonstrate comparable cellular binding or uptake. To differentiate between cell surface-bound and cell-internalized Aβ species (termed as ‘uptake’), we treated cells with trypsin-like cell dissociation reagent TrypLE™ Express (Thermo Fisher Scientific). TrypLE treatment removed surface-bound Aβ oligomers and fibrils while retaining intracellular puncta, which enabled us to distinguish astrocytic uptake vs. surface binding of Aβ oligomers and fibrils.

**Figure 2.**
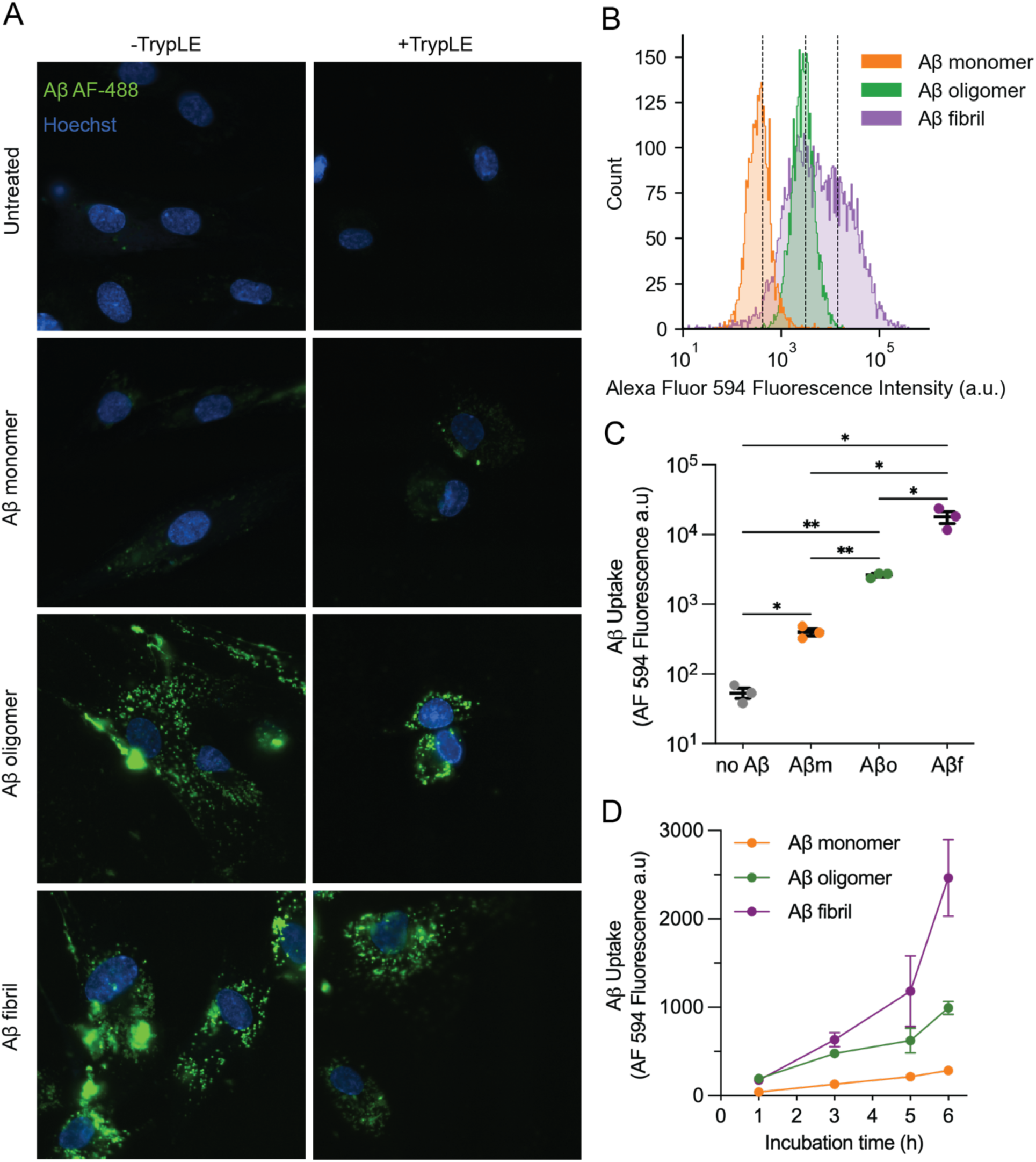
Aβ monomers, oligomers, and fibrils uptake by human astrocytes. **(A)** Live-cell fluorescence microscopy images showing the uptake of Alexa Fluor 488-labeled Aβ monomers, oligomers, and fibrils (1 µM total Aβ, with 7 nM labeled Aβ incorporated) by human cortical astrocytes after 24-hour incubation at 37°C. The “+TrypLE” panels show the cells that were treated with TrypLE to remove surface-bound Aβ, followed by an additional 24-hour recovery period. The “-TrypLE” panels show cells that were imaged immediately after 24-hour incubation. **(B)** A representative flow cytometry histogram showing the uptake of Alexa Fluor 594-labeled Aβ monomers, oligomers, and fibrils (18 nM labeled Aβ incorporated into 1 µM total Aβ) after 24 hours of incubation. **(C)** Quantification of Aβ uptake following 24 hours. The data are presented as individual biological replicates, with their mean ± SEM (n=3 biological replicates). Each biological replicate represents the average mean cell fluorescence from three technical replicates (at least 500 cells per replicate). Statistical significance was determined using Brown-Forsythe and Welch ANOVA test with post-hoc unpaired t-test with Welch’s correction, with individual variances computed for each comparison (*: P < 0.05; **: P < 0.01). (**D)** Aβ uptake as a function of incubation time over 6-hour period. Each data point represents the mean ± SD (n=3 technical replicates).

We further quantified Aβ uptake using flow cytometry, where fluorescence intensity in each cell served as a proxy for Aβ internalization. Similar to live cell imaging, minimal uptake was observed for Aβ monomers while higher uptake was observed for both oligomers and fibrils after 24-hour incubation (Fig. 2B). We also noted differences in the variance of the intensity distributions of uptake. Cells exposed to monomers and oligomers resulted in a relatively narrow cell intensity population, whereas cells exposed to Aβ fibrils resulted in a much wider range of fluorescence intensities, which indicated a higher heterogeneity of fibril uptake. This observation can also perhaps be attributed to the heterogeneity of fibril sizes reported earlier, where the uptake of distinct fibrils likely varies.

Quantitative comparison of the mean fluorescence intensities of the cells after 24-hour incubation further confirmed that the uptakes of monomers, oligomers, and fibrils were substantially different from each other (Fig. 2C). In fact, uptake was time-dependent for all aggregation states, continuously increasing over the observed time (Fig. 2D). Together, these observations suggest that the uptake of Aβ by astrocytes is significantly influenced by its aggregation state (fibril > oligomer > monomer), likely reflecting a protective mechanism where astrocytes preferentially clear aggregated Aβ species to buffer nearby neurons.

### Uniformity of ApoE monomers and lipoparticles is consistent across isoforms

The interaction between ApoE and Aβ is highly sensitive to the structural properties of both partners. Unlipidated ApoE is an aggregation-prone protein that exists in a concentration-dependent equilibrium of monomers, dimers, and tetramers^41^. ApoE4 was shown to have a higher propensity to self-associate with a lower tetramer-to-dimer and dimer-to-monomer dissociation constant compared to ApoE3 and ApoE2^41^. The propensity to self-associate could complicate biophysical measurements of complexation and increase the apparent dissociation constant (K_D_) of ApoE:Aβ complexes due to competition with ApoE self-association, similar to its impact on measuring ApoE-lipid binding^42^.

To deconvolve ApoE’s interaction with Aβ from competing self-association processes, we prepared ApoE at low nanomolar concentrations where monomeric species predominate^41^. We then used FCS to confirm that all three isoforms exist as monomers. Fitting of the FCS autocorrelation curve of ApoE with a one-component diffusion model yielded a uniformly reproducible R_H_ of approximately 4 nm across all three isoforms (Fig. 3B). This preparation and characterization allows us to attribute changes in diffusion purely to the interaction between ApoE monomers and Aβ monomers and aggregates.

**Figure 3.**
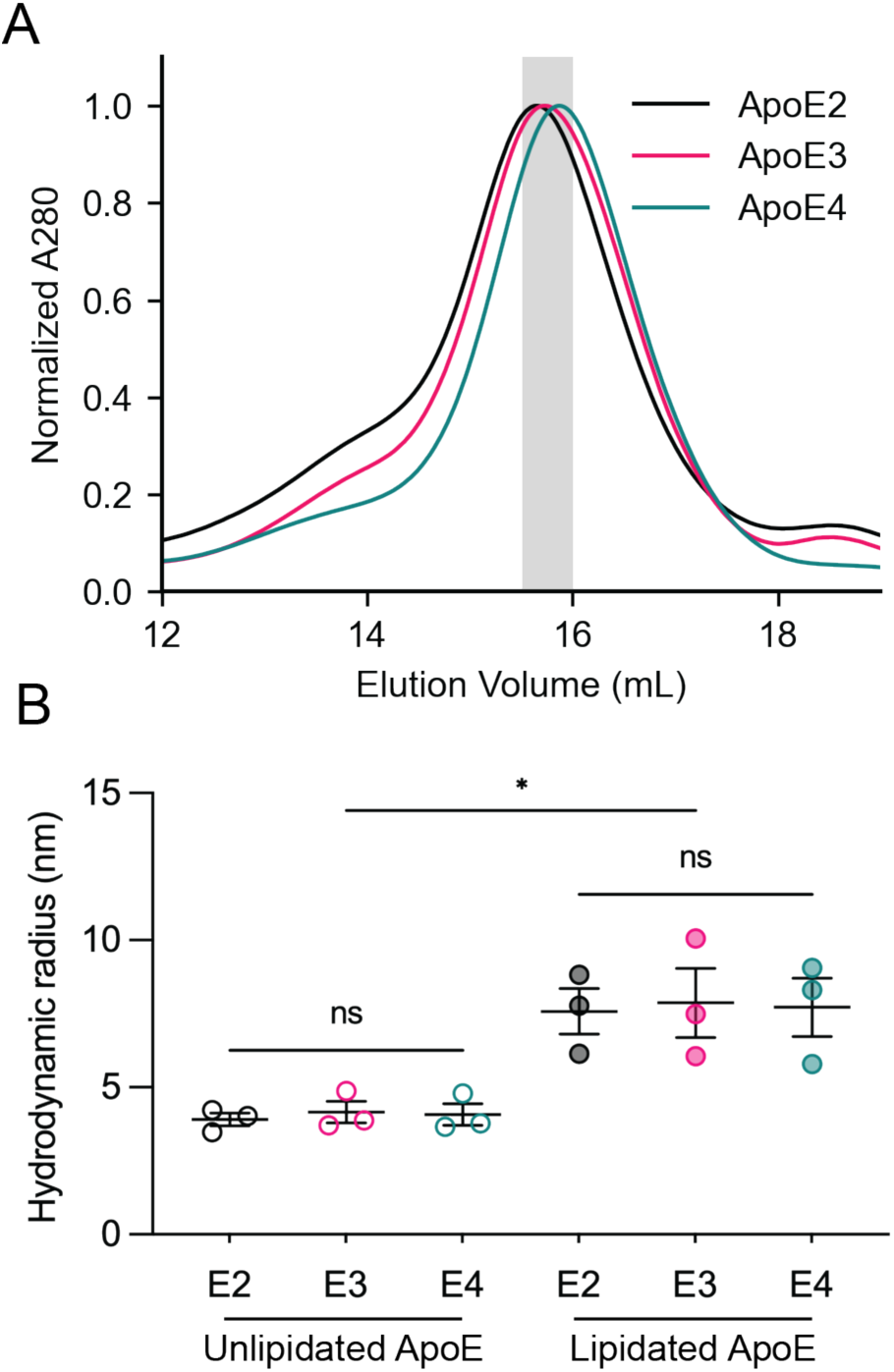
ApoE isoforms are equally lipidated. **(A)** Size exclusion chromatograms of ApoE isoforms lipidated with POPC and cholesterol. Absorbance at 280 nm is normalized to the highest absorbance value in each chromatogram. The gray shaded region indicates the elution fraction collected for subsequent analysis. **(B)** Hydrodynamic radii of 1 nM Alexa Fluor 488-labeled unlipidated and lipidated ApoE isoforms in PBS (pH 7.2). Values were derived from a single-component fit of the FCS autocorrelation curve. The results are expressed as individual protein and lipoparticle preparations, their mean ± SEM. Statistical significance was determined using ordinary one-way ANOVA with a post-hoc Tukey’s multiple comparisons test (*: P < 0.05, ns: P > 0.05).

We artificially lipidated ApoE with a mixture of POPC and cholesterol at a 1:90:5 ApoE:POPC:cholesterol ratio, which is standard in other studies and creates a stable reproducible reconstituted particle that closely resembles the size, morphology, and function of the High-Density Lipoprotein (HDL)-like particle in the brain^25,43^. Consistent with previous observations^44^, size exclusion chromatography (SEC) of prepared lipoparticles showed slight differences in the elution times between ApoE isoforms, further supporting the fact that the size of artificially lipidated ApoE is isoform isoform-dependent (ApoE2>ApoE3>ApoE4) (Fig. 3A).

To achieve our goal of decoupling isoform-specific differences in lipidation from the intrinsic sequence effects, we carefully controlled for lipoprotein particle size. We fractionated the main SEC elution peak of ApoE lipoparticles in 0.5 mL increments and selected an overlapping sub-fraction for all subsequent experiments (Fig. 3A, gray rectangle). We then used FCS to more precisely measure R_H_ on the selected fractions. We confirmed that ApoE was successfully lipidated by observing a significant increase in the R_H_ of ApoE lipoparticles compared to unlipidated ApoE (Fig. 3B). ApoE lipoparticles produced an FCS autocorrelation curve that was fit with a one-component diffusion model. Similar to the sizes of ApoE lipoprotein particles secreted by astrocytes^45,44^, the measured R_H_ values for isoforms were 7.6 ± 1.3 nm (ApoE2), 7.9 ± 2 nm (ApoE3), and 7.7 ± 1.7 nm (ApoE4). In other words, there were no significant differences in R_H_ between the lipidated isoforms, confirming that lipoparticles are equally lipidated. These observations also agree with previous studies of ApoE POPC lipoparticles with a diameter of 14 nm^46^.

### Unlipidated ApoE isoforms uniformly binds Aβ oligomers and fibrils, but not monomers

The physiological relevance of the interaction between ApoE and Aβ, particularly the relevance of certain Aβ aggregation states, remains a subject of debate^47^. To investigate if unlipidated ApoE selectively regulates distinct Aβ species, we characterized the binding affinity between unlipidated ApoE isoforms and Aβ prepared in distinct aggregation states. To do this we employed a fluorescence polarization (FP) assay, which reports on changes in the rotational diffusion of a fluorescently labeled molecule resulting from its interaction with a binding partner. In FP assays, optimal resolution is achieved by fluorescently labeling the smaller of two interacting species because the binding of a large, slow-moving molecule to a smaller one causes a significantly larger and more detectable decrease in rotational speed, which results in larger increases in FP signal^48^. For Aβ monomer binding to ApoE, we used SEC-purified FAM-labeled Aβ peptide (smaller species) (Fig. S1F) and kept its concentration at 1 nM, which is well below its reported critical aggregation concentration^37^. This ensured that we were specifically observing monomeric Aβ interactions. We then titrated increasing concentrations of unlipidated label-free ApoE (larger species) and measured the resulting changes in FP. Interestingly, we did not observe changes in FP upon addition of ApoE to Aβ monomers, indicating a lack of binding at the tested concentrations (100 nM – 10 μM) (Fig. 4A).

**Figure 4.**
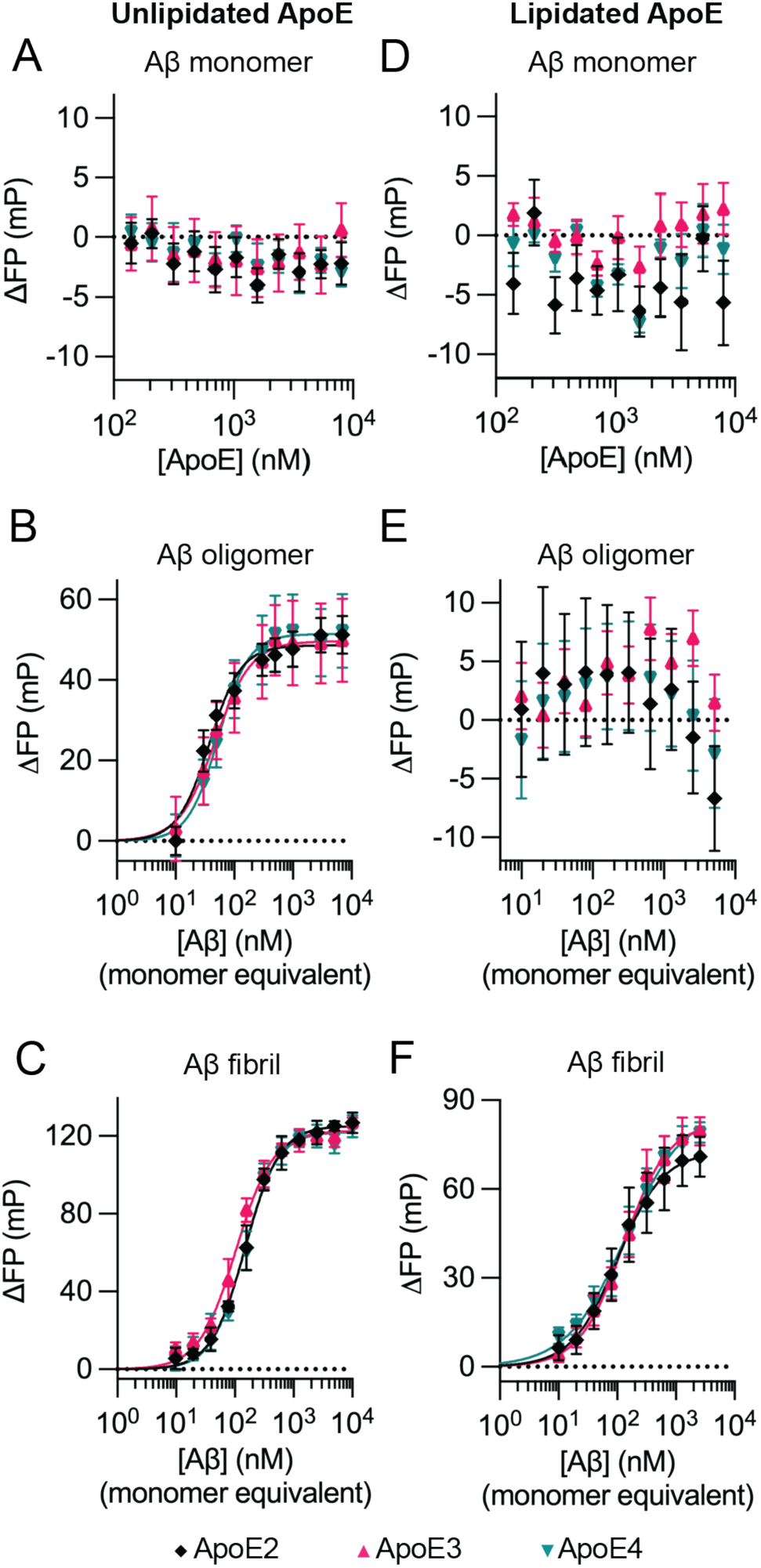
Fluorescence Polarization analysis of ApoE and Aβ interaction. Binding curves of 1 nM FAM-labeled Aβ incubated with 100 nM – 8 µM unlipidated ApoE **(A)** or lipidated ApoE **(D)**. Binding curves of 1 nM Alexa Fluor 488-labeled unlipidated ApoE incubated with 10 nM – 10 µM of pre-formed Aβ oligomers **(B)** or Aβ fibrils **(C)**. Binding curves of 1 nM Alexa Fluor 488-labeled lipidated ApoE incubated with 10 nM – 10 µM pre-formed Aβ oligomers **(E)** or fibrils **(F)**. The Y-axis represents normalized FP values, where the initial FP value of labeled Aβ monomer or labeled ApoE in the absence of titrated ligand is subtracted from each subsequent point. All samples were incubated for 2 hours at room temperature prior to measurement. Data represent the mean and SEM of triplicate measurements.

In order to measure the binding affinity of aggregated Aβ to ApoE, we used 1 nM of Alexa Fluor 488-labeled unlipidated ApoE (smaller species) and titrated increasing concentrations of label-free Aβ oligomers or fibrils (larger species). In both cases, we observed a concentration-dependent increase in FP when we added Aβ oligomers or fibrils to fluorescently labeled ApoE. We fit FP curves to a binding model with a Hill slope and obtained K_D_ values in the nM range, indicating very tight binding between unlipidated ApoE and Aβ oligomers and fibrils (Fig. 4B, C). The ability of unlipidated ApoE to bind Aβ oligomers and fibrils but not monomers, and a Hill slope greater than one, suggests a multivalent binding between ApoE and aggregated Aβ. We also observed a slightly higher binding affinity of unlipidated ApoE to Aβ oligomer over Aβ fibril (Table 1, Supplementary Table S2A), indicating that Aβ oligomers either have more ApoE binding sites, have binding sites with a higher affinity, or contain a more favorable spatial arrangement of binding sites necessary for multivalent interaction.

**Table 1.** Effect of ApoE isoform and lipidation on interaction with Aβ monomer, oligomer, and fibril. Dissociation constants were extracted using a saturation binding equation with a Hill Coefficient. Values represent the mean K_D_ ± SD (nM) from three measurements.

|  |  | Unlipidated ApoE |  | Lipidated ApoE |  |
| --- | --- | --- | --- | --- | --- |
| | | $K_D$ (nM) | $n_H$ | $K_D$ (nM) | $n_H$ |
| Aβ monomer | ApoE2 | >8 μM | N.D. | >8 μM | N.D. |
|  | ApoE3 |  |  |  |  |
|  | ApoE4 |  |  |  |  |
| Aβ oligomer | ApoE2 | 37 ± 2 | 1.6 ± 0.1 | >8 μM | N.D. |
| | ApoE3 | 45 ± 10 | 1.5 ± 0.5 | | |
| | ApoE4 | 53 ± 6 | 1.7 ± 0.1 | | |
| Aβ fibril | ApoE2 | 157 ± 39 | 1.6 ± 0.4 | 154 ± 135 | 1.2 ± 0.4 |
| | ApoE3 | 107 ± 24 | 1.4 ± 0.2 | 143 ± 67 | 1.2 ± 0.2 |
| | ApoE4 | 147 ± 36 | 1.7 ± 0.7 | 170 ± 98 | 0.9 ± 0.2 |

When comparing ApoE2, ApoE3, and ApoE4, we observed remarkably similar binding affinities to Aβ oligomers or fibrils (Table 1, Supplementary table S1A, S1B). This suggests that the direct interaction between ApoE and aggregated Aβ alone is not sufficient to reconcile the observed *in vivo* differences in how ApoE isoforms differentially impact Aβ clearance and deposition in AD.

### Lipidated ApoE binds Aβ fibrils, but not oligomers

We next surveyed how lipidated ApoE binds Aβ monomers, oligomers, and fibrils using a similar FP assay described above. Similar to unlipidated ApoE, we did not observe significant interactions between FAM-labeled Aβ monomers and unlabeled lipidated ApoE across the reported concentrations (Fig. 4D). Lipidated ApoE also retained its binding to Aβ fibrils with minimal differences from unlipidated ApoE (Fig. 4F, Supplementary Table S2B). Again, we did not observe large differences between isoforms (Table 1, Supplementary Table S1C). However, in contrast to unlipidated ApoE, we observed no binding between Alexa Fluor 488-labeled lipidated ApoE and unlabeled Aβ oligomers (Fig. 4E). This observation suggests that the Aβ oligomer binding sites on ApoE lipoparticle are occluded by the lipid, consistent with previous reports showing that Aβ competes with lipids for the binding sites on ApoE^28^.

To ensure that the lack of oligomeric binding is not a limitation of the FP assay, we performed a complementary FCS measurement, where we looked at changes in the diffusion of Alexa Fluor 488-labeled ApoE lipoparticles in the presence of Aβ oligomers. Similarly, we did not observe any changes to the R_H_ of lipidated ApoE when Aβ oligomers were added to all three ApoE isoforms, which confirms the lack of binding. In contrast, the FCS experiment conducted with unlipidated ApoE resulted in a significant increase in the average R_H_ of unlipidated ApoE when Aβ oligomers were added, which is indicative of binding. This observation agrees with the binding reported earlier using FP (Fig. S2). In fact, the average R_H_ of unlipidated ApoE:Aβ oligomer complex was significantly higher than the R_H_ of ApoE lipoparticles alone, which also suggests that FCS is sensitive enough to detect changes in ApoE lipoparticle diffusion when adding larger unlabeled Aβ oligomers.

In summary, the distinct binding patterns between lipidated and unlipidated ApoE highlight that the lipidation state of ApoE, not the isoform, critically modulates its interaction with Aβ. While other factors outside of the scope of these *in vitro* experiments likely supplement these discrepancies in AD pathology, it is notable that by controlling ApoE’s lipidation and aggregation state, the complexation between all ApoE isoforms (protective and pathological) and Aβ is identical. This implicitly suggests that mutations to ApoE indirectly tune its lipidation, which likely drives its regulatory behavior. This view is in contrast with the scenario where mutations to ApoE directly modulate the affinity to Aβ, thus driving the physiological differences in clearance.

### Unlipidated ApoE inhibits the astrocytic uptake of Aβ oligomers and fibrils, with no isoform differences

In addition to direct interactions, ApoE and Aβ both share similar cellular receptors ^20,22,49,50^. Therefore, we asked whether the presence of exogenous ApoE enhances Aβ uptake by astrocytes through receptor interactions or decreases Aβ uptake through competition for similar clearance pathways^20,43,47^. Both processes could also occur simultaneously, depending on local concentrations of free and complexed ApoE, which could explain contrasting observations^20,43^. To test this hypothesis we incubated immortalized human astrocytes with Alexa Fluor 594-labeled Aβ monomers, oligomers, or fibrils (1 μM) in the presence of different concentrations of unlipidated ApoE (1 – 128 nM). We then characterized how ApoE impacts Aβ uptake by comparing the intracellular intensity of fluorescently labeled Aβ in flow cytometry in the presence of, or absence of, exogenous ApoE.

Interestingly, we did not observe any enhanced uptake of Aβ in the presence of ApoE. Instead, our experiments revealed only a concentration-dependent inhibitory effect of unlipidated ApoE on astrocytic uptake of Aβ oligomers and fibrils (Fig. 5B, C), but not monomers (Fig. 5A). We fit these results to a one-phase exponential decay model to quantitatively compare the inhibitory effect between ApoE isoforms and aggregation states of Aβ (Table 2). Together, we find that unlipidated ApoE had the biggest effect on Aβ oligomers versus Aβ fibrils (Supplementary Table S4A), though similar to the binding studies we did not observe ApoE isoform impacting Aβ uptake (Supplementary Table S3A, B).

**Figure 5.**
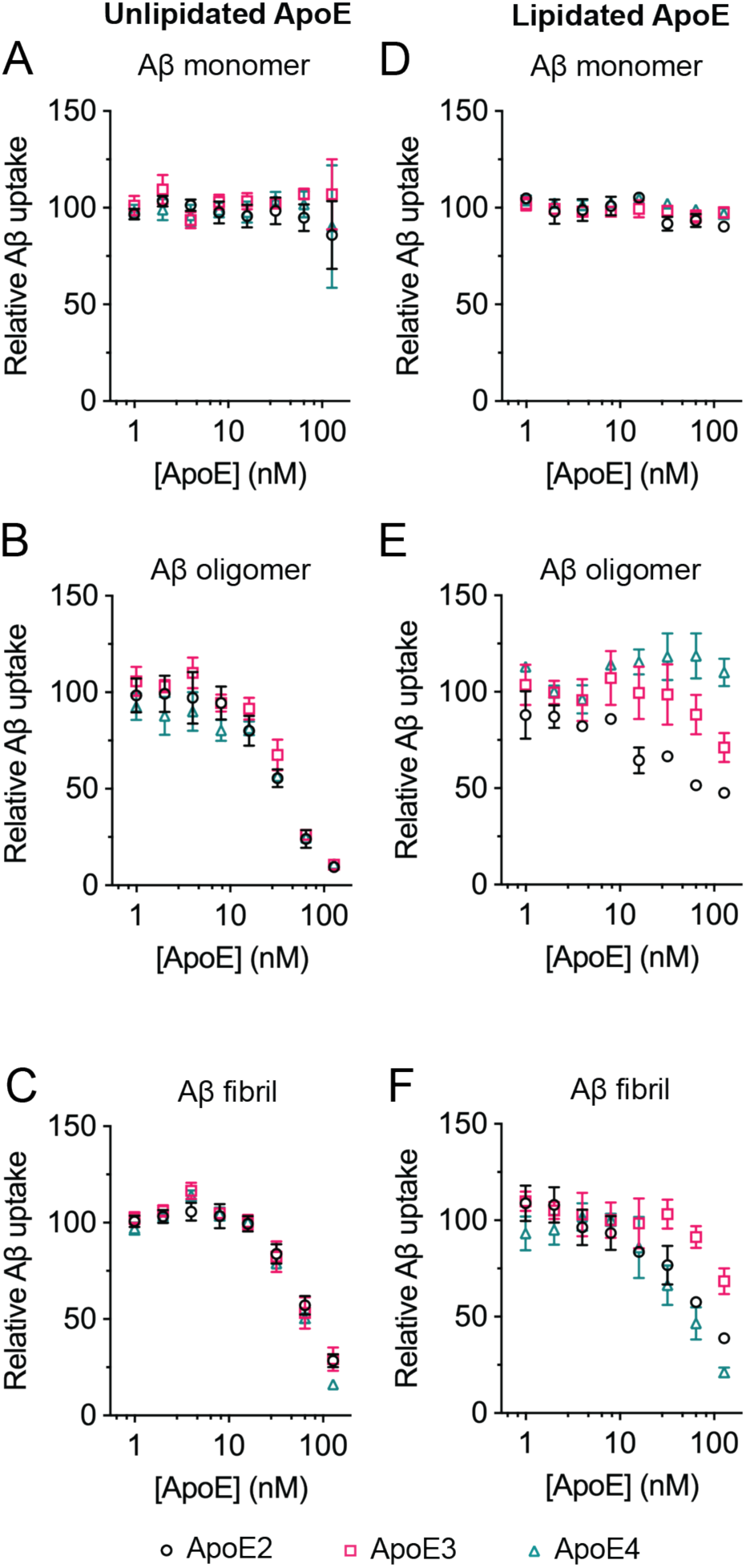
Effect of ApoE on Aβ uptake by astrocytes. Immortalized human cortical astrocytes were treated with 1µM Aβ monomers (A,D), oligomers (B,E), or fibrils (C,F), containing 18 nM Alexa Fluor 594 labeled species. Cells were incubated with or without varying concentrations (1 – 128 nM) of unlipidated ApoE **(A-C)** or lipidated ApoE **(D-F)** for 24 hours at 37°C. Aβ uptake, represented by intracellular Alexa Fluor 594 intensity, was measured via flow cytometry and normalized to Aβ uptake in the absence of ApoE. Results are expressed as mean ± SEM of three biological replicates. Each biological replicate is an average of three technical replicates, each containing at least 500 cells.

**Table 2.** Effect of ApoE isoform and lipidation on the uptake of Aβ monomer, oligomer, and fibril. Half-maximal inhibitory concentrations were extracted using the exponential decay equation to fit Aβ uptake as a function of ApoE concentration. Values represent the mean IC_50_ ± SD (nM) from three independent measurements.

| | $IC_{50}$ (nM) | | | | | |
| --- | --- | --- | --- | --- | --- | --- |
|  | unlipidated ApoE |  |  | lipidated ApoE |  |  |
| | Aβm | Aβo | Aβf | Aβm | Aβo | Aβf |
| ApoE2 | >128 nM | 35 ± 2 | 78 ± 16 | >128 nM | >128 nM<br>(E4>E3>E2) | 81 ± 7 |
| ApoE3 | | 40 ± 4 | 76 ± 24 | | | >128 nM |
| ApoE4 | | 39 ± 2 | 68 ± 6 | | | 60 ± 16 |

The inhibitory effect of unlipidated ApoE on Aβ oligomer and fibril uptake by protective astrocytes suggests a detrimental role of unlipidated ApoE in AD pathology. It is possible the unlipidated ApoE hinders the removal of aggregated Aβ from the extracellular space, further contributing to decreased clearance of toxic Aβ oligomers and increased deposition of Aβ fibrils. However, the lack of isoform differences in this study suggests that the inhibition of Aβ uptake alone cannot explain the isoform-specific differences of ApoE on disease pathology.

### Lipidated ApoE is less effective at inhibiting Aβ oligomer uptake, but results in comparable Aβ fibril uptake

Next, we investigated how different isoforms of equally lipidated ApoE lipoparticles influence Aβ monomer, oligomer, and fibril uptake by astrocytes. Similar to unlipidated ApoE, ApoE lipoparticles did not substantially modulate Aβ monomer uptake (Fig. 5D). For Aβ fibril uptake, ApoE lipoparticles had a similar inhibitory effect compared to unlipidated ApoE. Notably, ApoE3 lipoparticles demonstrated the least inhibitory effect on Aβ fibril uptake while ApoE2 and ApoE4 lipoparticles reduced Aβ uptake with similar efficacy as their lipid-free counterparts (Fig. 5F, Table 2, Supplementary Tables S3C).

However, a crucial difference emerged when we examined the effect of ApoE lipoparticles on Aβ oligomer uptake. ApoE lipoparticles generally had a smaller effect on Aβ oligomer uptake compared to unlipidated ApoE (Fig. 5E). Interestingly, this effect was isoform-dependent with ApoE2 lipoparticles exhibiting the highest reduction of Aβ oligomer uptake compared to ApoE3, while ApoE4 lipoparticles had no effect on Aβ oligomer uptake. These results align well with previously reported isoform-dependent differences in endocytosis of lipoprotein particles. ApoE2 lipoparticles are typically endocytosed less than ApoE3 and ApoE4^51^, allowing them to persist in media longer and inhibit Aβ oligomer uptake.

These observations initially seemed surprising, especially that protective ApoE2 lipoparticles inhibit the astrocytic uptake of Aβ oligomers. However, if we account for the inadequate lipidation of ApoE4 observed *in vivo*^26,27^ and compare unlipidated ApoE4 to lipidated ApoE2 and ApoE3, our results can be directly connected to the isoform-specific differences observed in AD pathology. Doing so suggests that poorly-lipidated ApoE4 would attenuate Aβ oligomer uptake and subsequent elimination of these toxic species by astrocytes (resulting in a buildup of toxic Aβ oligomer species) compared to lipidated ApoE3 and ApoE2.

### Aβ oligomers are toxic to human astrocytes, but toxicity can be reduced by lipidated ApoE

Finally, we assessed the effect of Aβ in different aggregation states on cell viability to check if there are functional consequences of our binding and uptake assays. We first performed a concentration-response experiment to determine a concentration of Aβ that induced significant cytotoxicity without causing complete cell death. We applied Aβ monomers, oligomers, and fibrils at 4 μM (moderate oligomer cytotoxicity) to immortalized human cortical astrocytes and measured cell viability after 24 hours using a Cell Titer Pro 2.0 luminescence reporter (Promega). We found significantly higher toxicity of Aβ oligomers to immortalized astrocytes compared to Aβ monomers and fibrils, which highlights a notable difference in their biological activity (Fig. 6, Fig. S3A). We then looked at the effect of ApoE on the toxicity of each Aβ aggregate. We treated cells with 4 μM of Aβ in different aggregation states in the presence (or absence) of lipidated and unlipidated ApoE (100 nM).

**Figure 6.**
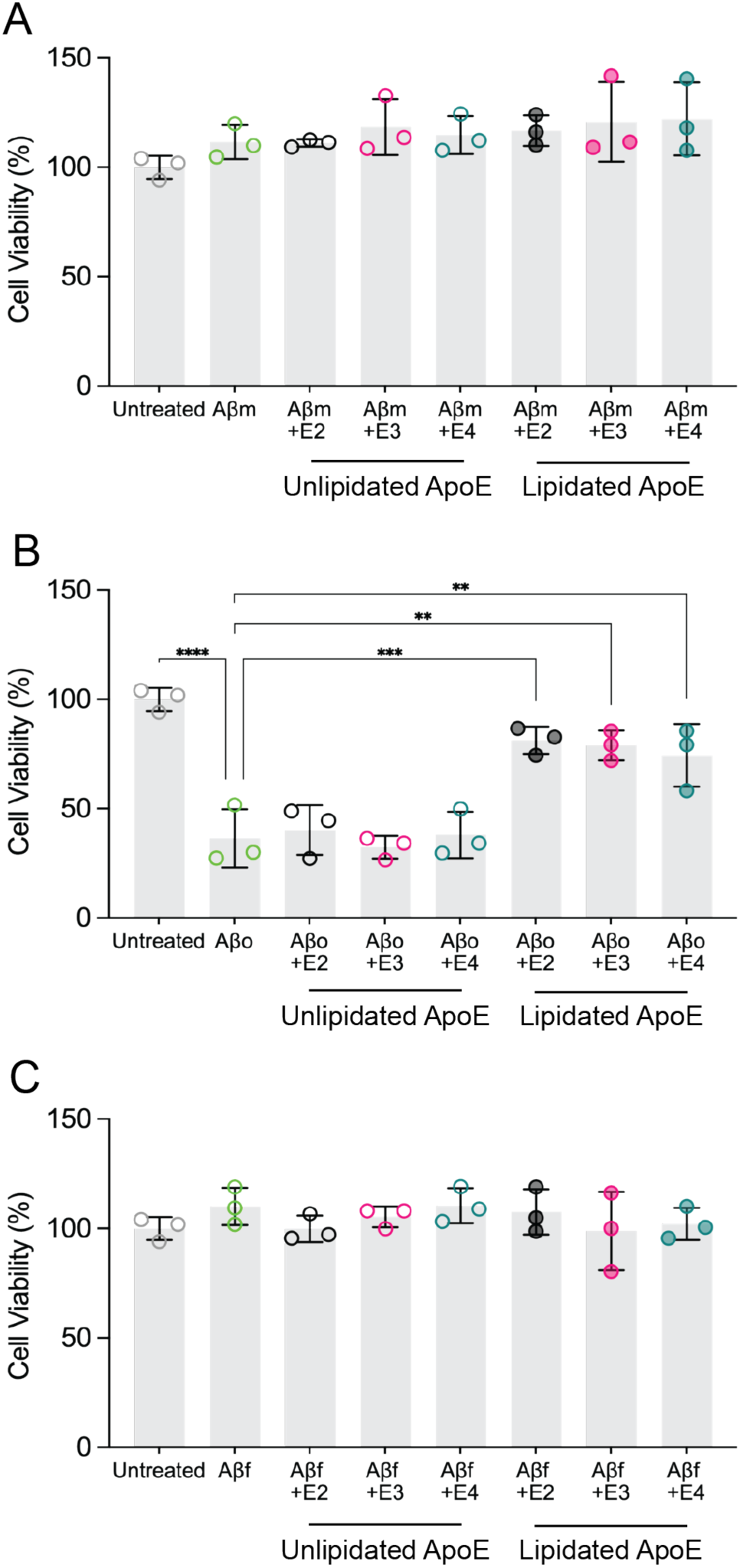
Effect of ApoE on Aβ-mediated cytotoxicity in astrocytes. Immortalized human cortical astrocytes were treated with 4 µM Aβ monomers (A), oligomers (B), or fibrils (C), containing 100 nM of unlipidated ApoE or lipidated ApoE for 24 hours at 37°C. Cell viability was measured as the number of live cells and normalized to vehicle-treated cells. Results are displayed as mean ± SD of three biological replicates, representing independent astrocyte cultures. Statistical significance was analyzed using Welch’s t-test, comparing each individual condition in the presence of ApoE to a respective Aβ monomer, oligomer, or fibril treatment in the absence of ApoE. Only significant differences are indicated on the graph (*: P < 0.05; **: P < 0.01, ***: P < 0.001, ****: P < 0.0001, ns: P > 0.05), all other comparisons are not significant.

Interestingly, the cytotoxicity of Aβ oligomers was independent of their uptake. Unlipidated ApoE, which previously reduced Aβ oligomer uptake, had no effect on its cytotoxicity (Fig. 6B). In contrast, lipidated ApoE, which had a much smaller effect on Aβ oligomer uptake, significantly decreased the toxicity of Aβ oligomers (Fig. 6B). This is consistent with our previous observations and suggests that unlipidated ApoE (largely ApoE4) does not rescue Aβ oligomer toxicity while lipidated ApoE (largely ApoE2 and ApoE3) therapeutically rescues Aβ oligomer toxicity. In the absence of Aβ oligomers, lipidated and unlipidated ApoE did not impact astrocyte viability on their own (Fig. S3B). Once again, the lipidation of ApoE, rather than the intrinsic differences in the effects of ApoE isoforms, appears to be the primary determinant that modulates Aβ cytotoxicity, which we believe has direct implications for AD pathology. Not only is ApoE4 less lipidated *in vivo*, but there are also lower expression levels of ApoE4 compared to other isoforms^52^. This suggest that therapeutic strategies to target ApoE-related pathologies should include modulators of ApoE4 lipidation in order to maintain normal astrocyte functioning through the sequestration and clearance of toxic Aβ oligomers.

## DISCUSSION

The observations in this study resolve long-standing mechanistic ambiguities regarding the role of ApoE in AD pathology. We demonstrate that the lipidation state of ApoE and the aggregation state of Aβ are the primary determinants of their interaction, subsequent uptake by astrocytes, and overall cytotoxicity. We achieved this detailed mechanistic understanding by normalizing ApoE lipoparticle size, Aβ aggregation state (monomers, oligomers, and fibrils) (Fig. 1), and by minimizing confounding variables such as isoform-specific self-association of ApoE (Fig. 3). This systematic quantitative comparison addresses the complex interplay between ApoE isoforms and their lipidation, and links these states to their putative effect on Aβ pathology.

Our findings challenge the notion that the intrinsic differences between ApoE isoforms directly modulate the affinity for Aβ, thereby driving physiological differences in clearance. Instead, we demonstrate that the lipidation of ApoE, rather than isoform, has the strongest effect on its interaction with Aβ (Fig. 4), on Aβ uptake by astrocytes (Fig. 5), and on Aβ cytotoxicity (Fig. 6). It is notable that when unlipidated ApoE isoforms were kept monomeric and ApoE lipoparticles kept equally lipidated (Fig. 3), the complexation between all isoforms of ApoE and Aβ was not significantly different. This insight explains many contradictory findings from the literature that report isoform differences in binding between ApoE and Aβ^53,54^. The studies often examine binding at concentrations where unlipidated ApoE self-associates, which increases the apparent dissociation constant between ApoE and Aβ. Our observations confirm that mutations to ApoE likely influence pathology indirectly through changes in lipidation, fundamentally contrasting the scenario where mutations directly affect Aβ association.

Furthermore, our work highlights the critical role of Aβ aggregation in its interaction with ApoE. We observed that toxic Aβ oligomers were distinctly affected by the ApoE lipidation state. Working with pre-formed purified oligomers allowed us to eliminate the confounding effects of monomers and fibrils, which typically vastly outnumber oligomers along the Aβ aggregation pathway^55^. While unlipidated ApoE strongly binds Aβ oligomers, lipidated ApoE loses this ability. In contrast, both lipidated and unlipidated ApoE bind Aβ fibrils while neither bind Aβ monomers, consistent with previous reports of multivalent interactions between ApoE and Aβ^21,56^. Measuring binding in solution using FP allowed us to circumvent key limitations of surface-based binding assays and avoid artifacts such as artificial avidity, multivalent interactions at a surface, and the potential for non-native Aβ conformations. Additionally, it enabled us to use Aβ monomers at very low concentrations, circumventing problems associated with the rapid aggregation of Aβ. Co-treatment with unlipidated ApoE resulted in more substantial inhibition of Aβ oligomer uptake by astrocytes compared to ApoE lipoparticles. Similar to the interaction studies, Aβ fibril uptake was equally modulated by both lipidated and unlipidated ApoE, while Aβ monomer uptake remained unaffected.

The cellular observations, combined with our biophysical data, suggest an alternative mechanism for ApoE-mediated reduction of Aβ uptake by human astrocytes distinct from competition for cellular endocytic receptors. Previous studies similarly observed decreased Aβ uptake in the presence of ApoE by different CNS cell types^20,57,22^, often attributing this to competition with cellular receptors and surface proteins such as LDLR, LRP1, and HSPG^20,22,49,50^. However, if receptor competition were the sole factor we would expect lipidated ApoE, which interacts more readily with endocytic receptors^58^, to be a more potent inhibitor of Aβ oligomer and fibril uptake compared to unlipidated ApoE. Our observations contradict this prediction. Instead, we propose that ApoE binding to Aβ oligomers (unlipidated ApoE only) and fibrils (both lipidated and unlipidated ApoE) reduce the accessibility of Aβ binding sites to cellular receptors and result in lower astrocyte uptake and clearance. This proposed mechanism is supported by existing therapeutic efforts to target the interaction between Aβ and ApoE, subsequently facilitating Aβ clearance^59^, which ameliorates disease progression in transgenic AD mice^60^.

By controlling for ApoE concentration and equal lipidation of ApoE isoforms, we were able to efficiently deconvolve the contributions of ApoE isoforms and lipidation on Aβ clearance by astrocytes. Contextualizing our results with physiological ApoE protein and lipidation levels in the brain further connects the effects observed in solution and *in vitro* experiments to actual AD pathology. In human iPSC-derived isogenic astrocytes^61^ and humanized ApoE mouse brains^52^ ApoE protein levels are genotype-specific (ApoE2 > ApoE3 > ApoE4) due to their stability and aggregation propensities, despite similarities in mRNA levels across the isoforms^52^. Moreover, the levels of lipidated ApoE and the amount of lipid per particle are also isoform-specific (ApoE2 > ApoE3 > ApoE4)^26,27^. This dual deficiency in both lipidation and ApoE4 levels creates a highly pathogenic environment where astrocytes are deficient in Aβ clearance and more sensitive to Aβ oligomer-induced stress. On the contrary, high levels of lipidated ApoE2 would help explain its protective role against AD.

Our data strongly agree with the emerging hypothesis that links the poor lipidation of ApoE to its pathogenicity, and reinforce the therapeutic strategy of targeting ApoE lipidation^24^. This link is strongly supported by genetic mutations to ApoE lipidation machinery such as ATP-binding cassette (ABC)-family transporters ABCA1 and ABCA7, which increase the risk of Alzheimer’s Disease^62^. The critical role of proper lipidation of ApoE is further supported by studies showing that the deletion of the ABCA7 transporter is associated with increased Aβ pathology in 5xFAD mice^63^. Additionally, in systems that mimic *in vivo* isoform-specific differences in ApoE levels and lipidation (such as humanized ApoE mice or iPSC-derived astrocytes), Aβ clearance and deposition are ApoE isoform-dependent (*APOE*-KO>*APOEε2*> *APOEε*3> *APOEε*4)^20,61^. Many promising interventions that modulate these transporters or ApoE-mediated lipid efflux such as LXR agonists, ApoE lipidation enhancers^64^, or antibodies targeting lipid-free ApoE^65^ demonstrate improved Aβ clearance and decreased Aβ accumulation. Our observations further strengthen this therapeutic route by providing detailed molecular mechanisms to explain how enhanced ApoE lipidation directly connects to improved Aβ clearance.

While our solution and *in vitro* experiments provide new mechanistic clarity through precise control over protein aggregation and lipidation conditions, they simplify the incredibly complex and dynamic brain environment, which contains multiple cell types, diverse lipoprotein populations, and heterogeneous mixtures of Aβ in co-existing aggregation states. Limitations to our study include the use of *E.coli*-derived (and artificially lipidated) ApoE that lack isoform-specific post-translational modifications and lipid diversity relevant to Aβ interactions^66^. While we used equally-lipidated ApoE lipoparticles to isolate the effect of lipidation, this normalization may mask subtle isoform differences stemming from intrinsic variations in lipoparticle size and stability. Future studies should build upon our findings and utilize astrocyte-derived ApoE particles to quantify the effect of ApoE on Aβ metabolism in multiple CNS cell types. Given the extensive literature on surface receptors involved in Aβ uptake such as LDLR, LRP1, and HSPG^20,22,49,50^, future studies should also focus on the role of specific receptors in ApoE-mediated inhibition of Aβ uptake using well-defined ApoE and Aβ species.

## CONCLUSION

The totality of our results emphasizes the critical roles of ApoE lipidation and Aβ aggregation on the interaction and subsequent uptake of Aβ by human astrocytes. We demonstrate that the effects of ApoE are specific to toxic Aβ oligomers, highlighting the necessity of using rigorously prepared species in future mechanistic studies. Moreover, our findings provide strong evidence that the *in vivo* protective or pathological effects of ApoE isoforms likely do not stem from intrinsic differences in how ApoE isoforms interact with Aβ, but rather from interstitial concentrations and varying degrees of lipidation. In this scenario, the protective role of ApoE2 can be best illustrated by its increased lipidation and expression levels, whereas the risk associated with ApoE4 is driven by poor lipidation. These findings have important implications for mitigating the increased Alzheimer’s Disease risk associated with ApoE4. Our data support the therapeutic potential of both enhancing ApoE4 lipidation and inhibiting the interaction between unlipidated ApoE and toxic Aβ oligomers. We demonstrate that therapeutic efforts to mitigate ApoE4-related pathologies should be extremely careful to account for Aβ aggregation state and the biochemical properties of ApoE, such as its lipidation and aggregation state, to maximize the power of *in vitro* experiments for interpreting complex *in vivo* phenomena.

## MATERIALS AND METHODS

### Aβ oligomer preparation

Aβ oligomers were prepared according to a previously published protocol^34^, with modifications to accommodate recombinantly expressed peptide and the presence of NaOH. Reagent volumes were calculated to achieve a final Aβ concentration of 1.67 mg/mL (364 μM) in 0.2% SDS in 1X PBS (pH 7.4) and to compensate for 50 mM NaOH in Aβ stock. The initial incubation volume, typically 300 μL, was achieved by combining concentrated Aβ solution in 50 mM NaOH containing 1 mg of peptide, 10X low salt PBS (81 mM Na_2_HPO_4_, 15 mM KH_2_PO_4_, 870 mM NaCl, 27 mM KCl), 10X PBS (81 mM Na_2_HPO_4_, 15 mM KH_2_PO_4_, 1370 mM NaCl, 27 mM KCl) (Thermo Fisher Scientific, J62036.K7), 10% SDS (Invitrogen, 24-730-020), 1 M HCl, and MilliQ water. Reagents were added in this specific order to prevent rapid pH fluctuations that could cause undesired peptide aggregation. After mixing, the sample was incubated at 37°C for 5 hours. After the incubation, the solution was diluted with 3x volume (900 μL) of MilliQ water and incubated for an additional 18 hours at 37°C. The sample was then centrifuged at 20,000 x *g* for 20 minutes and concentrated 10-fold using a 30 kDa MWCO Amicon Ultra-0.5 centrifugal filter (Millipore, UFC5030) (120 μL final volume). The concentrated oligomers were dialyzed using Slide-A-Lyzer™ MINI Dialysis Devices, 20K MWCO (Thermo Fisher Scientific, 88405) against 0.25X PBS (pH 7.4), with two changes of buffer (2 hours, 2 hours, 18 hours), centrifuged at 20,000 x *g* for 20 minutes, aliquoted, and stored in -80°C until use.

### Aβ fibril preparation

Aβ fibrils were prepared as previously described^35^. Briefly, purified recombinant Aβ monomer in 50 mM NaOH was concentrated to approximately 150 μM. Peptide solution was then neutralized with 1 M HCl. Additional volumes of 1 M HCl and MilliQ were then added to achieve 100 μM Aβ solution in 10 mM HCl. Aβ was incubated for 48 hours at 37°C with rigorous shaking. The solution was then rigorously vortexed to resuspend the formed fibrils. Fibrils were subsequently aliquoted and stored at -80°C. To prepare the sonicated Aβ fibril, the fibril solution in 10 mM HCl was first neutralized with 50 mM NaOH and resuspended to the desired concentration and volume in PBS. This solution was then sonicated in a bath sonicator (Fisherbrand, FB11201) at 37 kHz and 70% amplitude for 10 minutes in ice-cold water. The sonicated fibrils were used immediately. To prepare fluorescently labeled Aβ oligomers and fibrils, fluorescently labeled Aβ monomers were mixed with unlabeled Aβ monomers at desired ratios before the aggregation reactions.

### FCS instrumentation and setup

All FCS experiments were conducted using a MicroTime 200 STED time-resolved fluorescence microscope (PicoQuant, GbmH, Berlin, Germany), equipped with 60X water-immersion objective with correction collar (NA 1.2, UPlanSApo, Olympus) and fitted with an NKT Photonics SuperK FIANUM White Light Laser (WLL) (440–790 nm). A pulsed 485 nm excitation laser was used for Alexa Fluor 488 excitation, while a NKT Photonics SuperK VARIA tunable filter was used to select 590 nm excitation for Alexa 594 from WLL emission. The WLL was operated at a pulse repetition rate of 20 MHz, and the SYNC output of WLL was used to trigger interleaved pulses from the 488 diode laser. The average laser power was 2.5 μW (measured at the back aperture of the objective) for both excitation wavelengths to minimize fluorophore triplet state conversion.

Samples (200 μL) were placed into 8-well Bioinert μ-Slides (IBIDI, 80800). Lasers were focused to approximately 20 μm above the coverslip. The system was calibrated using Rhodamine 110 and free Alexa Fluor 594 dyes, referencing previously published diffusion coefficients^67,68^. For each measurement, 6-10 traces, each 30 seconds in duration, were recorded. Autocorrelation curves were calculated using the SymPhoTime 64 software package (PicoQuant, GbmH, Berlin, Germany). Fluorescence lifetime correlation spectroscopy (FLCS) was used to remove background and after-pulsing, and time-gating was used to separate Alexa Fluor 488 and Alexa Fluor 594 excitation periods (PIE-FCS).

Each 30-second trace was fit individually using a 3D diffusion model (single species (n=1 in the equation below) for Aβ monomers, ApoE monomers, or ApoE lipoparticles; two species (n=2) for aggregated Aβ or binding studies) and a triplet term. For Aβ oligomers and fibrils, the diffusion time of monomeric Aβ was a fixed term in the two-species fit.

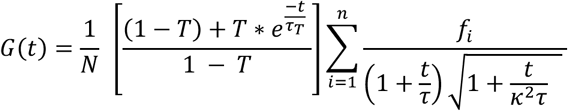

Here, G(t) is the autocorrelation function as a function of time (t). *τ* is the translational diffusion time of labeled molecules, N is the average number of fluorescent species in the confocal volume, *κ*^2^ is the ratio of axial to radial dimensions of the confocal volume, and f_i_ is the fraction of each of the components (SUM(f_i_) = 1). T is the average fraction of molecules in the triplet state, and *τ*_T_ is the triplet state lifetime.

### FCS sample preparation

Alexa Fluor 594-labeled Aβ monomers were initially prepared at 5 μM in 50 mM NaOH, then diluted 1000-fold with PBS (pH 7.2) to 5 nM. For the studies in cell media, 5 nM Alexa Fluor 594-labeled Aβ monomers were mixed with unlabeled 1μM Aβ monomers in phenol red-free, fetal bovine serum (FBS) free Astrocyte medium (ScienCell, 1801-PRF). Alexa Fluor 594-labeled Aβ oligomers (1:56 and 1:200 labeling ratios) and sonicated fibrils (1:1000, 1:2000, and 1:10000 labeling ratios) were diluted to 3 μM (total Aβ) in PBS (pH 7.2). 200 μL of sample was measured at room temperature in 8-well Bioinert μ-Slides (IBIDI, Cat. No. 80800).

For the studies looking at the interaction between ApoE and Aβ, Alexa Fluor 488-labeled ApoE or ApoE/POPC/cholesterol particles were diluted to 1nM in the μ-slide well. ApoE was allowed to dissociate to its monomeric form for at least 2 hours prior to the addition of Aβ. Subsequently, 3 μM Aβ oligomers were added to ApoE and incubated for at least 2 hours at room temperature and then measured.

### Cell Culture

Human cortical astrocytes (ScienCell, 1800) were seeded at 5,000 cells/cm^2^ and cultured as adherent monolayers in T-75 or T-175 tissue culture flasks at 37 °C in a 5% CO_2_ humidified incubator. The cells were maintained in Astrocyte Medium, supplemented with 5% fetal bovine serum (FBS), astrocyte growth supplement, and penicillin/streptomycin solution (ScienCell, 1801). Cells were typically used within 2-8 passages and routinely tested for mycoplasma contamination. A stable line of human cortical astrocytes was generated via transduction of human cortical astrocytes (ScienCell, 1800) with hTERT lentivirus. Human cortical astrocytes, immortalized with hTERT, were cultured in the exact same way as human cortical astrocytes. Cells retained their morphology and ability to divide as late as passage 22.

### Widefield fluorescence live cell imaging of Aβ uptake

Primary human cortical astrocytes were plated at a 5,000 cells/well density on 96-well flat-bottom polystyrene tissue culture-treated plate (Corning, 3997) and cultured for 24 hours prior to treatments. Alexa Fluor 488-labeled Aβ monomers, oligomers, and fibrils were pre-mixed at 7 μM in ice-cold PBS (pH 7.2) in Protein LoBind 1.5 mL tubes (Eppendorf). The solutions were then immediately diluted 7-fold to a final concentration of 1 μM with FBS-free Astrocyte Medium to prevent aggregation of Aβ monomers. The cell culture medium was removed from the astrocytes, and 100 μL of Aβ-containing solutions were applied to each well. Cells were incubated with proteins for 24 hours.

Following incubation, cells were either imaged immediately to assess both surface-bound and internalized Aβ, or treated with a trypsin-like reagent TrypLE Express (Gibco, 12604013) to remove surface-bound Aβ. For the latter, cell media containing Aβ monomer, oligomer, or fibrils was removed, cells were washed twice with 200 μL of Dulbecco’s phosphate-buffered saline (DPBS) (Gibco, 14190144), and then treated with 60 μL of TrypLE. After a 10-minute incubation at 37°C, TrypLE activity was quenched with 140 μL of fresh Astrocyte Medium. Cells were then transferred to a new 96-well glass-bottom plate with high-performance #1.5 cover glass (Cellvis, P96-1.5H-N) and centrifuged for 5 min at 300 x *g* at 4°C. The supernatant was removed, and the cells were resuspended in 100 μL of Astrocyte Medium. Cells were allowed to re-adhere for 24 hours before subsequent imaging.

All live-cell imaging was performed on an ImageXpress Confocal HT.ai High-Content Imaging System (Molecular Devices), utilizing a 40X water immersion objective. Cell nuclei were stained with Hoechst 33342 (Invitrogen, H3570) at 1 μg/mL for 10 minutes prior to the measurement. All imaging was performed at 37°C and 5% CO_2_ in Phenol Red-Free Astrocyte Medium (ScienCell, 1801-PRF). Images were analyzed using IN Carta Software (Molecular Devices).

### Flow cytometry of Aβ uptake

Immortalized human astrocytes were plated at 5,000 cells/well on a 96-well flat-bottom polystyrene tissue culture-treated plate (Corning, 3997). Cells were incubated for 24 hours prior to the start of the experiment to ensure recovery of cell surface receptors. Alexa Fluor 594-labeled Aβ monomer, oligomer, and fibrils were prepared with a labeling ratio of 1:56 (labeled to unlabeled Aβ). All samples were first mixed at a total concentration of 7 μM (total Aβ) in PBS and diluted sevenfold with FBS-free astrocyte medium to a final concentration of 1 μM. Cells were treated with 100 μL of this protein solution for 24 hours. For the time course uptake experiment, cells were plated at the same time and at the same density and treated with 1 μM Aβ monomers, oligomers, and fibrils 1, 3, 4, 5, or 6 hours prior to a single simultaneous measurement.

For experiments investigating the effect of ApoE, cells were treated with the same Alexa Fluor 594-labeled Aβ monomers, oligomers, and fibrils (18nM labeled Aβ incorporated into 1 μM total Aβ) in the presence of SEC-purified lipidated and unlipidated ApoE at concentrations ranging between 1 nM and 128 nM for 24 hours. ApoE was prepared at different concentrations via serial dilutions in 1.5 mL Protein LoBind tubes (Eppendorf). ApoE and Aβ were initially mixed in PBS (pH 7.2) and then diluted sevenfold with FBS-free astrocyte medium. To maximize complex formation, Aβ oligomer and fibril samples were incubated with ApoE for at least two hours prior to dilution with cell medium. Aβ monomer-containing samples were diluted immediately to avoid peptide aggregation in PBS. Cells were treated with 100 μL of the protein solution containing ApoE and Aβ for 24 hours. After the incubation, cells were washed once with DPBS, treated with TrypLE to detach cells and remove the surface-bound Aβ, and pelleted by centrifugation. Cells were then resuspended in cold PBS, containing 5% FBS.

Aβ uptake was quantified by measuring Alexa Fluor 594 fluorescence of individual astrocytes via fluorescence-activated cell sorting. At least 500 cells were quantified per condition using BD FACSymphony™ A3 Cell Analyzer (BD Biosciences) equipped with a high-throughput sampler. Cells were gated based on forward and side scatter to identify cell-like events and single-cell signals. The same gates were used for all experiments. Data was analyzed and visualized using FlowJo 10.10.0, Python, and GraphPad Prism. Data was normalized to the uptake of Aβ in the absence of ApoE (100%). The average fluorescence intensity of cells in the absence of Aβ was used as a baseline (0%).

### Fluorescence polarization to measure binding between FAM – Aβ monomers and unlabeled ApoE

FAM-labeled Aβ (BACHEM, 409015) was dissolved in 50 mM NaOH at 1 mg/mL and purified in 50 mM NaOH using a Superdex 200 Increase 10/300 GL column (Cytiva) to ensure that all of the aggregates were removed (Fig. S1F). SEC-purified peptide was first diluted to 5 μM in 50 mM NaOH, followed by a rapid 1000x dilution to 5 nM in 1x PBS (pH 7.2). 5 nM Aβ was then centrifuged for 20 minutes at 21,000 x *g* at 4°C to remove any aggregates that might have formed during the dilution process.

For binding titration, unlipidated ApoE and lipidated ApoE were serially diluted in a 96-well half-area black flat-bottom polystyrene non-binding surface microplate (Corning, 3993). To 40 μL of different dilutions of ApoE, 10 μL of 5 nM FAM-Aβ in PBS was added. To correct for the background fluorescence, a separate set of wells, containing unlabeled unlipidated ApoE and lipidated ApoE dilutions, was incubated without FAM-Aβ. All the samples were incubated for 2 hours at room temperature prior to measurement.

Fluorescence Polarization was measured using an EnVision 2105 Multimodal Plate Reader, equipped with FITC FP mirror, FITC FP 480 excitation filter, and two emission filters: FITC FP P-pol 535, and FITC FP S-pol 535 (Perkin Elmer). Each well was read 3 times, and the average was used for calculation.

Fluorescence polarization of FAM-Aβ was calculated in Python using the following formula:

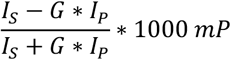

Here I_s_ is the fluorescence intensity of emitted light from a parallel direction in relation to the excitation plane, and I_p_ is the fluorescence intensity of emitted light from a perpendicular direction in relation to the excitation plane. Background fluorescence, measured from samples without FAM-Aβ, was subtracted from both I_s_ and I_p_. G is G-factor, a correction factor accounting for differences in instrument sensitivity when measuring I_s_ and I_p_. The G-factor for the instrument was determined using free Alexa Fluor 488 dye in PBS as a calibrant (FP of 28 mP).

### Fluorescence polarization to measure binding between Alexa Fluor 488-labeled ApoE and unlabeled Aβ oligomers and fibrils

Unlipidated Trp210Cys ApoE mutants, labeled with Alexa Fluor 488, were diluted to 2 nM in PBS and allowed to dissociate from the concentrated stock for 2 hours to ensure that ApoE is in its monomeric form. Label-free Aβ oligomer and fibril were serially diluted (10 nM to 10 μM final concentrations) in a 96-well half-area black flat-bottom polystyrene non-binding surface microplate (Corning, 3993). Equal volumes (20 μL) of unlipidated ApoE or ApoE/POPC/Cholesterol labeled with Alexa Fluor 488 (1 nM final concentration) were mixed with equal volumes (20 μL) of Aβ oligomers and fibrils in PBS (pH 7.2) in the 96-well microplate. To measure background fluorescence, unlabeled Aβ oligomers and Aβ fibrils were incubated without Alexa Fluor 488-labeled ApoE. FP values were calculated as described above.

For direct binding, the changes in FP values compared to the free ligand were plotted against the concentration of added protein. K_D_ was determined from fitting FP data in GraphPad Prism using a one-site specific binding with Hill Coefficient model.

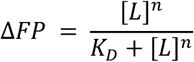

Here, ΔFP is a difference in FP between ApoE bound to Aβ oligomer or Aβ fibrils and free ApoE, [L] is Aβ oligomers or Aβ fibrils concentration, K_D_ is the dissociation constant, and n is the Hill Coefficient.

### Cellular toxicity of Aβ

Immortalized human cortical astrocytes were plated at 5,000 cells/well in 96-well glass-bottom plate with high-performance #1.5 cover glass (Cellvis, P96-1.5H-N) and allowed to adhere for 24 hours. 4 μM Aβ monomer, oligomer, and fibril samples were mixed with and without 100 nM ApoE or ApoE/POPC/Cholesterol in FBS-free astrocyte medium. 100 μL of pre-mixed protein solution was applied to each well for 24 hours. After incubation, 100 μL of Cell Titer Glo 2.0 reagent (Promega, G9242) was added to each well and incubated following the manufacturer’s recommendations. The luminescence signal in each well was measured using a EnVision 2105 Multimodal Plate Reader (Perkin Elmer). Data were normalized to cells that did not undergo any Aβ treatment.

## Abbreviations

AD: Alzheimer’s disease
Aβ: Amyloid Beta
ApoE: Apolipoprotein E
CSF: cerebrospinal fluid
FCS: fluorescence correlation spectroscopy
R_H_: hydrodynamic radius
FP: fluorescence polarization
FCCS: Fluorescence Cross Correlation Spectroscopy
ABC: ATP-binding cassette
POPC: 1-palmitoyl-2-oleoyl-sn-glycero-3-phosphocholine
PBS: phosphate-buffered saline
DPBS: Dulbecco’s phosphate-buffered saline

