## Supplemental Information for "ApoE lipidation, not isoform, is the key modulator of Aβ interaction, uptake, and cytotoxicity"

### SUPPLEMENTARY FIGURES

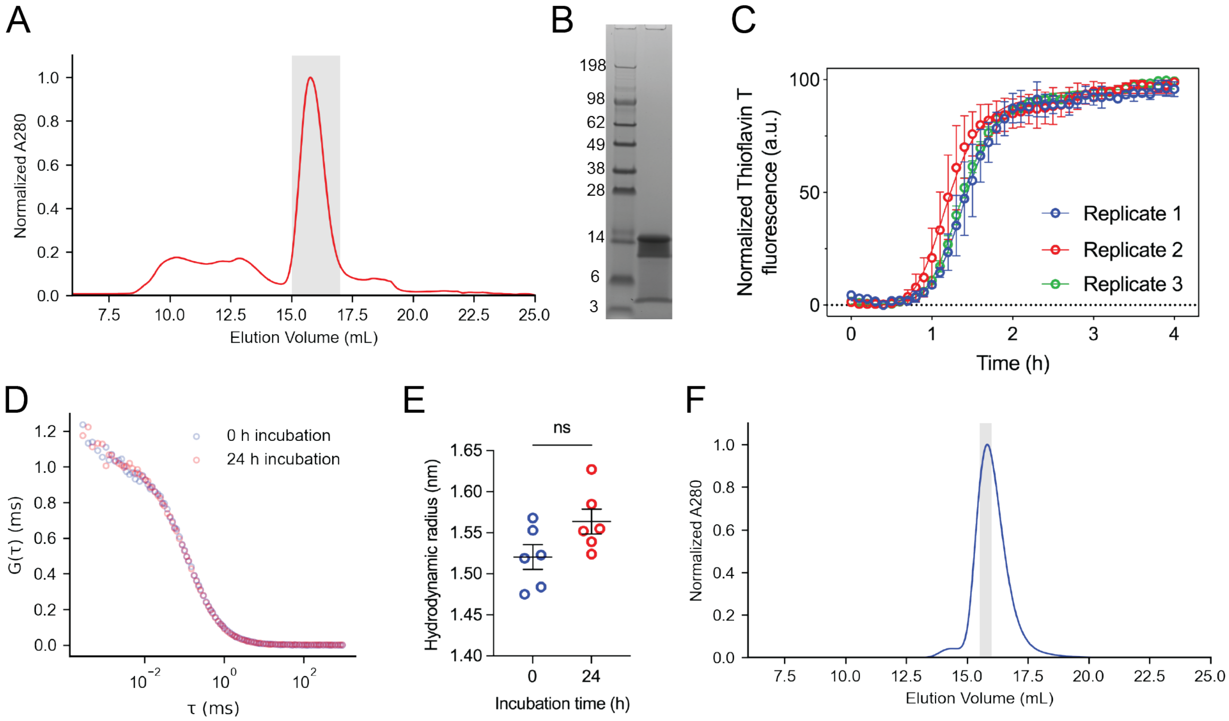


**Supplementary Figure 1. Preparation of pure aggregate-free Aβ monomer solutions. (A)** Size exclusion chromatogram of Aβ monomers eluted with 50 mM NaOH. The major peak (15 – 17 mL) represents Aβ monomers eluted with 50 mM NaOH. Minor peaks (7.5 – 15 mL) represent aggregated peptide and impurities. The gray shaded region indicates the collected fraction. **(B)** SDS-PAGE of Aβ peptide confirming its high purity. **(C)** ThT aggregation assay of 2.5 μM Aβ in 1x PBS (pH 7.4), normalized to maximum fluorescence at 4 hours of incubation, shows consistent aggregation of peptide, confirming aggregate-free starting material. **(D)** Overlapping FCS autocorrelation curves of Alexa 594-labeled Aβ monomers (~ 5 nM) in the presence of 1 μM unlabeled Aβ monomers confirm the stability of Aβ monomers in FBS-free Astrocyte Media at 37 °C over 24 hours. (**E)** Hydrodynamic Radii derived from a 3D diffusion model fit of FCS autocorrelation curves at 0 and 24 hours of incubation. Markers represent individual measurements of the sample (n=6). Error bars represent the mean ± SEM. Statistical significance was calculated using an unpaired t-test (ns: $P > 0.05$). **(F)** Size exclusion chromatogram of FAM-labeled Aβ monomers eluted with 50 mM NaOH, showing a narrow main peak consistent with monomer elution. The gray shaded region indicates the fraction collected for FAM-Aβ and ApoE interaction experiments.

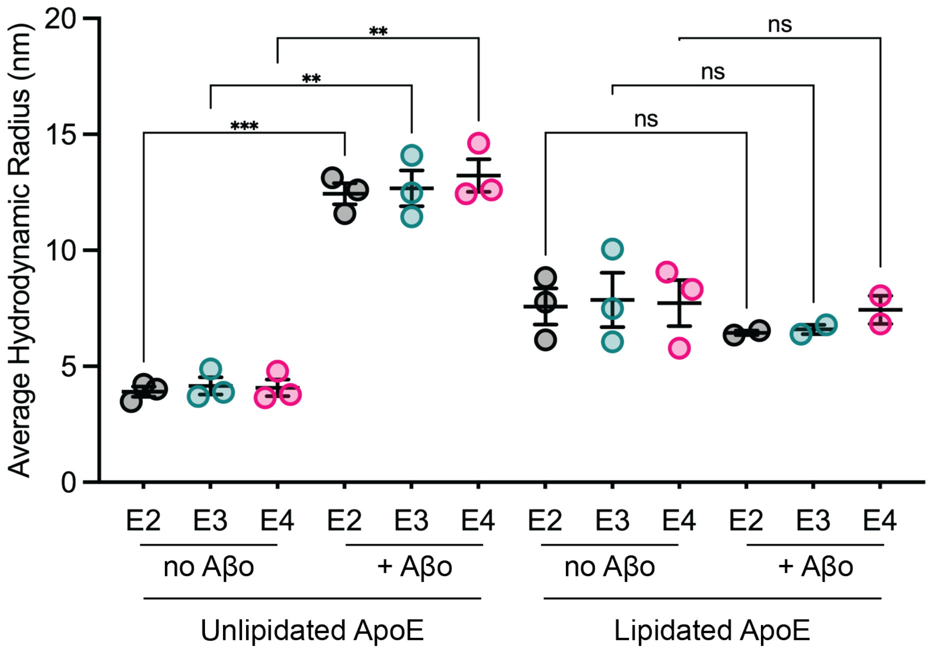


**Supplementary Figure 2. Hydrodynamic radii of unlipidated and lipidated ApoE in the presence and absence of Aβ oligomers.** The average hydrodynamic radii of 1 nM Alexa Fluor 488-labeled unlipidated and lipidated ApoE isoforms were measured after incubation with or without 3 μM pre-formed Aβ oligomers in PBS (pH 7.2). If lipidated ApoE were to bind Aβo, the average hydrodynamic radii should be greater than or equal to that observed for unlipidated ApoE + Aβo. The results are expressed as mean ± SEM. Each data point represents a measurement from an independent Aβ oligomer preparation. Statistical significance was determined using Brown-Forsythe and Welch ANOVA test with post-hoc unpaired t-test with Welch's correction, with individual variances computed for each comparison (**: $P < 0.01$, ***: $P < 0.001$, ns: $P > 0.05$).

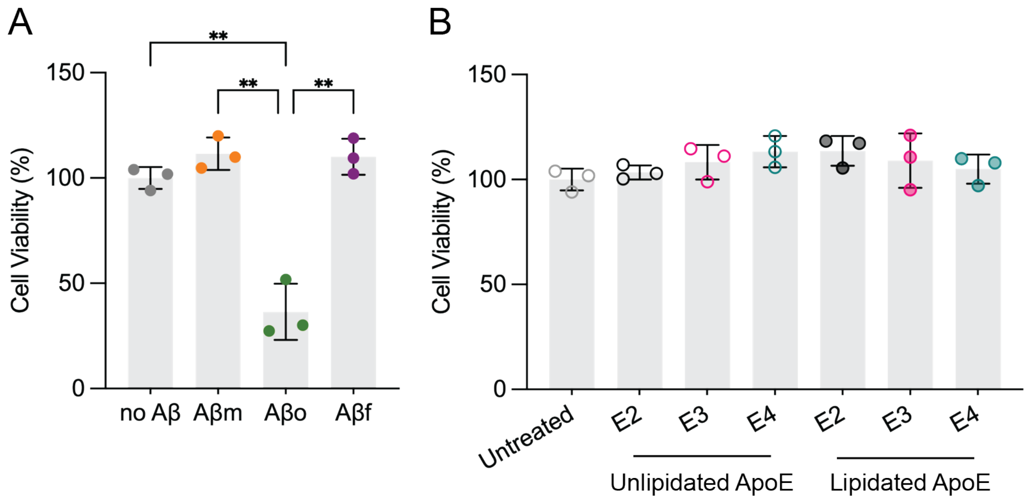


**Supplementary Figure 3. Cytotoxicity of Aβ monomers, oligomers, and fibrils and ApoE to immortalized human astrocytes. (A)** Aβ aggregation state influences its toxicity to immortalized human astrocytes. Cells were treated with 4 μM Aβ monomers, oligomers, or fibrils for 24 hours at 37°C. **(B)** Neither lipidated nor unlipidated ApoE (100 nM) by itself alters immortalized astrocyte cell viability. Cells were treated with 100 nM lipidated and lipid-free ApoE for 24 hours at 37°C. Cell viability, measured as the number of live cells, was normalized to vehicle-treated cells. Results are displayed as mean ± SD of three biological replicates, representing independent astrocyte cultures. Statistical significance was analyzed using Brown-Forsythe and Welch ANOVA test with post-hoc unpaired t-test with Welch's correction, with individual variances computed for each comparison (*: $P < 0.05$; **: $P < 0.01$). The effect of ApoE was compared to the untreated sample. Non-significant differences were observed.

### SUPPLEMENTARY TABLES

**Table S1.** The statistical significance of the differences in binding between ApoE isoforms and Aβ ($K_D$ values), determined by Tukey's multiple comparison test.

(A) Unlipidated ApoE and Aβ oligomer binding (Related to Fig. 4B, Table 1)

| | Summary | Individual P Value |
|---|---|---|
| ApoE2 vs. ApoE3 | ns | 0.4704 |
| ApoE2 vs. ApoE4 | ns | 0.0626 |
| ApoE3 vs. ApoE4 | ns | 0.3020 |

(B) Unlipidated ApoE and Aβ fibril binding (Related to Fig. 4C, Table 1)

| | Summary | Individual P Value |
|---|---|---|
| ApoE2 vs. ApoE3 | ns | 0.2427 |
| ApoE2 vs. ApoE4 | ns | 0.9288 |
| ApoE3 vs. ApoE4 | ns | 0.3773 |

(C) Lipidated ApoE and Aβ fibril binding (Related to Fig. 4F, Table 1)

| | Summary | Individual P Value |
|---|---|---|
| ApoE2 vs. ApoE3 | ns | 0.9889 |
| ApoE2 vs. ApoE4 | ns | 0.9831 |
| ApoE3 vs. ApoE4 | ns | 0.9462 |

**Table S2.** The statistical significance of the differences in binding between ApoE lipidation states and Aβ ($K_D$ values) grouped by ApoE isoform and determined by t-test with Welch's correction

(A) Unlipidated ApoE binding to Aβ oligomer vs. Aβ fibril (Related to Fig. 4B and 4C)

| | Summary | Individual P Value |
|---|---|---|
| ApoE2 | ** | 0.0073 |
| ApoE3 | * | 0.0305 |
| ApoE4 | * | 0.0438 |

(B) Unlipidated ApoE vs. lipidated ApoE binding to Aβ fibril (Related to Fig. 4C and 4F)

| | Summary | Individual P Value |
|---|---|---|
| ApoE2 | ns | 0.9802 |
| ApoE3 | ns | 0.4616 |
| ApoE4 | ns | 0.7356 |

**Table S3.** The statistical significance of the differences between ApoE isoforms in the inhibition of Aβ uptake, as measured by half-maximum ($IC_{50}$ values), was determined by Tukey's multiple comparison test.

(A) Aβ oligomer uptake inhibition by unlipidated ApoE (Related to Fig. 5B, Table 2)

| | Summary | Individual P Value |
|---|---|---|
| ApoE2 vs. ApoE3 | ns | 0.1097 |
| ApoE2 vs. ApoE4 | ns | 0.2413 |
| ApoE3 vs. ApoE4 | ns | 0.8140 |

(B) Aβ fibril uptake inhibition by unlipidated ApoE (Related to Fig. 5C, Table 2)

| | Summary | Individual P Value |
|---|---|---|
| ApoE2 vs. ApoE3 | ns | 0.9878 |
| ApoE2 vs. ApoE4 | ns | 0.7313 |
| ApoE3 vs. ApoE4 | ns | 0.8126 |

(C) Aβ fibril uptake inhibition by lipidated ApoE (Related to Fig. 5F, Table 2)

| | Summary | Individual P Value |
|---|---|---|
| ApoE2 vs. ApoE3 | ** | 0.0073 |
| ApoE2 vs. ApoE4 | ns | 0.8545 |
| ApoE3 vs. ApoE4 | ** | 0.0043 |

**Table S4.** The statistical significance of the differences between ApoE lipidation states in Aβ uptake inhibition grouped by ApoE isoform and determined by t-test with Welch's correction

(A) Aβ oligomer vs. Aβ fibril uptake inhibition by unlipidated ApoE (Related to Fig. 5B and 5C)

| | Summary | Individual P Value |
|---|---|---|
| ApoE2 | ** | 0.0091 |
| ApoE3 | ns | 0.0608 |
| ApoE4 | ** | 0.0014 |

(B) Aβ fibril uptake inhibition by unlipidated vs. lipidated ApoE (Related to Fig. 4B and 4C)

| | Summary | Individual P Value |
|---|---|---|
| ApoE2 | ns | 0.8173 |
| ApoE3 | * | 0.0171 |
| ApoE4 | ns | 0.4599 |

### SUPPLEMENTARY MATERIALS AND METHODS

#### Aβ42 expression and purification

The Aβ42 peptide, with methionine as its N-terminal residue (Aβ(Met 1-42)), was expressed in *Escherichia coli* (*E.coli*) following a published protocol[1]. In brief, the Aβ(Met 1-42) coding plasmid, which included an ampicillin resistance sequence (GenScript), was transformed into One Shot™ BL21 Star™ (DE3) chemically competent *E.coli* cells (Thermo Fisher Scientific, C601003).

Following the expression, the peptide was purified from inclusion bodies that were solubilized in 8 M urea using a bulk ion-exchange chromatography protocol with DEAE cellulose (Santa Cruz Biotechnology, sc-211213). Further purification was performed via size exclusion chromatography (SEC) using Superdex 200 Increase, 10/300 GL column (Cytiva) at 4°C (Fig. S1A). A 50 mM NaOH solution was used as an elution buffer to prevent peptide aggregation due to high Aβ elution concentrations[2]. Purified Aβ peptide was collected from the center of the elution peak and was either used immediately or flash-frozen in liquid $N_2$ and stored at -80°C. The purity of the peptide was confirmed via sodium dodecyl sulfate-polyacrylamide gel electrophoresis (SDS-PAGE). Purified Aβ was loaded on a Novex™ Tricine Mini Protein Gels, 10 to 20% (Invitrogen, EC6625BOX) and stained with SimplyBlue™ Coomassie stain (Invitrogen, LC6060) (Fig. S1B). To minimize peptide loss, Protein LoBind 1.5 mL tubes (Eppendorf) were exclusively used throughout all experiments involving Aβ.

**Aβ fluorescent labeling**

Purified Aβ monomer in 50 mM NaOH was diluted to 180 μM with ice-cold PBS (pH 7.4). The pH of the solution was then adjusted to 7.8 to optimize labeling efficiency. Dilution and pH adjustment steps were performed on ice as fast as possible to prevent Aβ aggregation prior to labeling. Alexa Fluor 594 succinimidyl ester (Thermo Fisher Scientific, A20004) was added to the Aβ solution at a 1:4 protein-to-dye molar ratio. The mixture was incubated for 2 hours in a Protein LoBind 1.5 mL tube (Eppendorf) at room temperature with continuous tumbling. After the 2-hour incubation, the reaction was quenched with 1 M Tris-HCl (Thermo Fisher Scientific, J22638-K2) (100 mM final concentration) for 10 minutes. To dissolve any Aβ aggregates that may have formed during incubation, 1 M NaOH was added to a final concentration of 250 mM. The solution was then sonicated in a bath sonicator, containing ice-cold water at 37 kHz and 50% amplitude for 10 minutes. The solution was subsequently centrifuged at 21,000 x *g* for 10 minutes at 4°C. The supernatant was then filtered using 0.22 μm Costar Spin-X centrifuge tube filters (Corning, 8160). Finally, the labeled peptide was SEC-purified using a Superdex 200 Increase, 10/300 GL column (Cytiva) to separate labeled protein from free dye, using 50 mM NaOH as an elution buffer. Eluted samples were immediately aliquoted into Protein LoBind tubes (Eppendorf), snap frozen in liquid nitrogen, and stored at -80°C.

**Thioflavin T assay**

Thioflavin T aggregation kinetic experiment was performed as previously described[3]. Aβ peptide in 50 mM NaOH was diluted to 2.5 μM in PBS (pH 7.2) and supplemented with 20 μM ThT from a 1 mM stock in 1.5 mL Protein LoBind Tube (Eppendorf) on ice. Samples were transferred into a half-area clear flat-bottom polystyrene non-binding surface microplate (Corning, 3881). The samples were covered with a plastic film (Biorad, MSB1001B). ThT fluorescence was measured through the bottom of the plate using an EnVision 2105 Multimodal Plate Reader, equipped with CFP 430 excitation and CFP 486 emission filters (Perkin Elmer). Measurements were collected at room temperature every 6 minutes with 5 minutes of shaking at 100 rpm in between measurements.

**ApoE Trp210Cys construct expression and purification**

To enable site-specific maleimide labeling of ApoE, necessary for FCS and FP assays, tryptophan at position 210 was mutated to cysteine (Trp210Cys) in all three ApoE isoforms. Maleimide-based labeling was chosen over amine labeling to ensure high labeling specificity and to avoid labeling numerous functionally important amine-containing lysine residues in ApoE. Trp210 is not a part of predicted structural elements, has low sequence conservation, and high predicted solvent accessibility. To prevent labeling of native cysteines, Cys112 and Cys158 in ApoE2, and Cys112 in ApoE3 were mutated to serines (Cys112Ser and Cys158Ser).

The ApoE Trp210Cys mutant coding plasmids (GenSript) were transformed into One Shot BL21 Star™ (DE3) chemically competent *E.coli* (Thermo Fisher, C601003). The ApoE was expressed as the fusion protein containing Thioredoxin (Trx) tag connected to 6x-His tag, separated from the ApoE sequence by a TEV protease cleavage site.

Protein was expressed in 1 L culture of *E.Coli* in LB Broth (IPM Scientific, 11006-010). Cells were harvested by centrifugation at 4,000 x *g* for 15 minutes. The cell pellets were lysed in 40 mL 1x Tris-buffered saline (TBS) (pH 8.0, 0.5 M NaCl) (BioRad, 1706435), containing 10% glycerol (Thermo Scientific, A16205.0F), 2 mM DTT (Goldbio, DTT100), 1 complete EDTA free Protease inhibitor (Roche, 04693132001), 0.1 mg/mL Lysozyme (Sigma, L6876), 0.01 mg/mL DNAse(Sigma, 04536282001), 0.01mg/mL RNAse (Roche, 10109134001). The lysate was then sonicated on an ice bath for 5 minutes (5 seconds on, 15 seconds off) cycles, at 50% amplitude using a Q500 immersion tip sonicator (Qsonica). The lysate was centrifuged at 20,000 x *g* for 30 minutes at 4 °C.

The supernatant was applied to nickel-nitriloacetic (Ni-NTA) resin (Thermo Scientific, 88223) and allowed to bind for 30 minutes. The resin was then washed with TBS supplemented with 30mM imidazole and 4 M guanidinium chloride (GdmCl) (Sigma-Aldrich, G3272) to remove impurities bound to ApoE. The fusion protein was then eluted using 300 mM imidazole (Thermo Scientific, A10221.36) in 1x TBS, containing 3.6 M GdmCl.

The eluted fusion protein was dialyzed using a 10 kDa MWCO dialysis cassette (Thermo Scientific, 87725) against 1x TBS, 1 mM DTT at room temperature, with three buffer changes over a 2-hour interval to remove GdmCl. To cleave the fusion protein, 1000 units of TEV (Sigma, T4455) were added to each dialysis cassette. Protein was cleaved overnight at room temperature with an additional change of dialysis buffer (1x TBS, 1 mM DTT) after 2 hours.

Following the cleavage, solid GdmCl was added to the ApoE solution to a final 4 M concentration. Solution was applied to a Ni-NTA column to remove the Trx-tag. ApoE was collected as a flow-through and was either immediately purified via SEC by Superdex 200 Increase, 10/300 GL column (Cytiva) in PBS (pH 7.2) or frozen in liquid $N_2$ and stored at -80°C.

To prevent the aggregation of protein, protein concentration was kept below 10 μM. Unlipidated ApoE readily binds to plastic, significantly decreasing the observed concentration over time. To prevent loss of protein to plastic adsorption, ApoE was kept in Protein LoBind tubes (Eppendorf). Additionally, protein concentration was measured prior to each experiment. When measuring concentrations of ApoE via A280 absorbance using NanoDrop Microvolume spectrophotometer (Thermo Scientific), the plastic pipette tip was primed with ApoE by repeated pipetting of protein solution (10-15 times) prior to the measurement. Aliquots of pure ApoE were stored at -80°C. To ensure reproducibility, stocks were not repeatedly freeze-thawed and were used within 5 days of thawing.

**Fluorescent labeling of ApoE**

Three isoforms of ApoE were labeled at Cys210 with Alexa Fluor 488 (Invitrogen, A10254) or Alexa Fluor 594 (Invitrogen, A10256) maleimide dyes following a published labeling protocol[4]. In brief, prior to labeling reaction, proteins were pre-treated with 10 mM DTT for 30 minutes to reduce cysteines and remove any potentially formed disulfide bonds. Proteins were then buffer exchanged into a DTT-free buffer containing 4 M GdmCl in 1x PBS (pH 7.4) using PD-10 desalting columns packed with Sephadex G-25 resin (Cytiva, 17085101) to remove DTT.

ApoE was incubated with a 5-10x molar excess of the maleimide dye for 2 hours at room temperature, protected from the light. The reaction was quenched with 10 mM DTT. Free dye was separated from labeled protein via SEC, using the Superdex 200 Increase, 10/300 GL column (Cytiva) in 1x PBS (pH 7.2)

**Apolipoprotein E lipidation**

ApoE was lipidated using the cholate dialysis method as previously described . POPC (Avanti Lipids, 850457C) and Cholesterol (Avanti Lipids, 700100) were dissolved in chloroform at 10 mg/mL. POPC and cholesterol were mixed at 18:1 molar ratio in a glass vial with a Teflon cap. 0.5 mL of the lipid mixture in chloroform was dried under nitrogen stream until a thin lipid film was formed. The lipid film was further dried under vacuum overnight. Lipid film was rehydrated with 1 mL 1x PBS (pH 7.2) for 30 minutes (final POPC concentration of 4.74 mM). After that, the lipid mixture was Vortexed vigorously for 2 minutes, 3 times with 2-minute intervals. Sodium Cholate (50 mg/mL solution in MilliQ) (Sigma-Aldrich, C6445-100G) was slowly added to the lipid solution until it was no longer cloudy. Concentrated lipid solution was added to 5-10 μM ApoE in 1x PBS at 1:90:5 ratios (ApoE:POPC:cholesterol). Sodium cholate was removed via dialysis against PBS (pH 7.2) using a 3.5 kDa MWCO dialysis cassette (Thermo

Scientific, A52966) over 36 hours at room temperature with several changes of buffer. Formed lipoparticles were concentrated using a 50 kDa MWCO Amicon 4 concentrator filter (Millipore, UFC 8050) and purified via SEC, using Superose 6 Increase, 10/300 GL column (Cytiva). To prepare fluorescently labeled ApoE lipoparticles, fluorescently labeled ApoE was used and the same protocol was followed.
